# Sensory neurons couple arousal and foraging decisions in *C. elegans*

**DOI:** 10.1101/2023.04.14.536929

**Authors:** Elias Scheer, Cornelia I. Bargmann

## Abstract

Foraging animals optimize feeding decisions by adjusting both common and rare behavioral patterns. Here, we characterize the relationship between an animal’s arousal state and a rare decision to leave a patch of bacterial food. Using long-term tracking and behavioral state classification, we find that food leaving decisions in *C. elegans* are coupled to arousal states across multiple timescales. Leaving emerges probabilistically over minutes from the high arousal roaming state, but is suppressed during the low arousal dwelling state. Immediately before leaving, animals have a brief acceleration in speed that appears as a characteristic signature of this behavioral motif. Neuromodulatory mutants and optogenetic manipulations that increase roaming have a coupled increase in leaving rates, and similarly acute manipulations that inhibit feeding induce both roaming and leaving. By contrast, inactivating a set of chemosensory neurons that depend on the cGMP-gated transduction channel TAX-4 uncouples roaming and leaving dynamics. In addition, *tax-4-*expressing sensory neurons promote lawn-leaving behaviors that are elicited by feeding inhibition. Our results indicate that sensory neurons responsive to both internal and external cues play an integrative role in arousal and foraging decisions.

## INTRODUCTION

Persistent behavioral states subdivide continuous behavior into discrete modules that accomplish adaptive goals (Tinbergen, 1951). An example from behavioral ecology is foraging behavior, which is composed of locomotion patterns that unfold across short and long timescales. For example, the brief darting maneuvers that drive prey capture in hunting zebrafish are embedded within persistent locomotory arousal states that last for minutes (Marques et al., 2020). In recent years, classical ethological studies of behavioral states have been supplemented with machine vision, clustering, and classification algorithms that are well-suited to quantitative and systematic analysis (Berman et al., 2014; Schwarz et al., 2015; Wiltschko et al., 2015). A current challenge is to identify circuit mechanisms that couple long-term behavioral states to short-term motor actions in foraging and other goal-directed behaviors.

Foraging behaviors and the neural mechanisms that generate them have been a fruitful subject of study in the nematode *Caenorhabditis elegans.* While exploring a bacterial lawn, which corresponds to a food patch, *C. elegans* adopts two major behavioral states: roaming and dwelling (Ben Arous et al., 2009; Fujiwara et al., 2002). Roaming is an aroused state characterized by high locomotion speed with few reversals and turns, whereas dwelling is defined by low speed and high reorientation rates (Ben Arous et al., 2009; Flavell et al., 2013; Fujiwara et al., 2002). A third behavioral state, quiescence, is induced by stress, molting, or satiety (Gallagher et al., 2013; Hill et al., 2014; Raizen et al., 2008; Van Buskirk & Sternberg, 2007). The bistability of roaming and dwelling states is governed by neuromodulatory signaling via the neuropeptide Pigment Dispersing Factor (PDF-1), which promotes roaming by signaling through its cognate receptor PDFR-1, and the biogenic amine serotonin, which promotes dwelling by activating the serotonin-gated chloride channel MOD-1. These modulators act through distributed circuits within the 302 neurons of the animal’s nervous system, with multiple sources of each modulator and multiple receptor-expressing sites (Flavell et al., 2013).

Another foraging behavior studied in *C. elegans* is the decision to stay or leave a lawn of bacterial food. Leaving is infrequent on high quality food lawns but increases when feeding is impaired, the bacterial food is inedible, or food is depleted (Bendesky et al., 2011; Milward et al., 2011; Olofsson, 2014; Shtonda and Avery, 2006). Animals also leave more frequently from lawns of pathogenic bacteria (Pujol et al., 2001) and lawns spiked with aversive or toxic compounds (Melo & Ruvkun, 2012; Pradel et al., 2007).

The genes that regulate lawn-leaving behavior overlap with those that regulate roaming and dwelling states, although these behaviors have largely been studied separately. For example, roaming, dwelling, and leaving are all strongly influenced by sensory neurons, particularly those that express the cGMP-gated channel *tax-4* (Davis et al., 2023; Fujiwara et al., 2002; McCloskey et al., 2017; Milward et al., 2011) (Table 1), and by endocrine DAF-7 (TGF-beta) and Insulin/DAF-2 (insulin receptor) signaling pathways whose peptide ligands are produced by sensory neurons (Ben Arous et al., 2009; Bendesky et al., 2011).

**Table 1:** Sensory neurons and endocrine systems implicated in roaming and leaving behavior*.

|  |  |  |  |  |
| --- | --- | --- | --- | --- |
| **Condition** | **Sensory/**<br>**endocrine system** | **Effect** | **Evidence/mechanism** | **Citation** |
| OP50 bacterial lawn | ciliated sensory neurons | Increase roaming | *che-2* cilium structure mutant | Fujiwara 2002 |
|  | ciliated sensory neurons | Increase roaming | rescue of *grk-2* mutant | Davis et al 2023 |
|  | *tax-4*-expressing neurons | Increase roaming | rescue of *che-2* mutant | Fujiwara 2002 |
|  | AWC neurons | Increase roaming | *ceh-36* mutant | Ben Arous 2009 |
|  | ASI neurons | Increase roaming | Tetanus toxin expression | Greene 2016 |
|  | BAG neurons | Increase roaming | Caspase induced ablation | Juozaityte 2017 |
|  | ASG neurons | Increase roaming | Caspase induced ablation | Juozaityte 2017 |
|  | ASJ neurons (male) | Increase leaving | rescue of *daf-7* TGF-beta mutant | Hilbert 2017 |
|  | *daf-2* insulin receptor | Increase roaming | mutant | Ben Arous 2009 |
|  | *daf-7* TGF-beta | Increase roaming | mutant | Ben Arous 2009 |
| OP50, icas#9 pheromone | ASJ neurons | Increase roaming | pheromone response, tetanus toxin expression | Greene 2016 |
|  | ADL neurons | Reduce roaming | pheromone response, tetanus toxin expression | Greene 2016 |
| OP50, depleting lawn | BAG neurons | Increase leaving | rescue of *tax-2* mutant, ablation | Milward 2011 |
|  | ASE neurons | Increase leaving | rescue of *tax-2* mutant, ablation | Milward 2011 |
|  | URX/AQR/PQR neurons | Increase leaving | rescue of *tax-2* mutant, ablation | Milward 2011 |
|  | AFD neurons | Increase leaving | rescue of *tax-2* mutant | Milward 2011 |
|  | ADF neurons | Reduce leaving | rescue of *ocr-2* mutant | Milward 2011 |
|  | ASI neurons | Increase leaving | rescue of *daf-7* TGF-beta mutant | Milward 2011 |
|  | *daf-2* insulin receptor | Increase leaving | mutant | Milward 2011 |
|  | *daf-7* TGF-beta | Increase leaving | mutant | Milward 2011 |
| Related behaviors (OP50) | ASI neurons | Increase quiescence | ablation | Gallagher 2013 |
| \**che-2*, *ceh-36*, *tax-2*, *tax-4*, and *ocr-2* mutations result in reduced sensory neuron function.<br>Studies in *npr-1* and *egl-4* mutants, which have an increased leaving rate, are not included because they have complex effects on neuronal activity.<br>Results addressing leaving from lawns of pathogenic bacteria were not included. |  |  |  |  |

Here, we characterize the behavioral patterns and circuit mechanisms that accompany lawn leaving decisions on short and long timescales. Using high-resolution imaging to monitor the behaviors of many individual animals, we find that leaving is a discrete behavioral motif that is generated probabilistically during the high-arousal roaming state. A signature of leaving behavior is a rapid acceleration in speed immediately before the leaving event. Roaming states evoked by neuromodulatory mutations, acute feeding inhibition, or optogenetic circuit manipulation all stimulate lawn leaving accompanied by the rapid acceleration motif. In addition, chemosensory neurons play an unexpectedly central role in linking internal states and behavioral dynamics.

## RESULTS

### Animals make foraging decisions at the boundaries of bacterial food lawns

To study foraging decisions at high resolution, we filmed and quantified the behavior of individual animals on small lawns of bacterial food (∼3 mm diameter) on an agar surface (Fig. 1A,B). During a 40-minute assay, animals spent ∼97% of their time on the lawn, and preferentially occupied the edge of the bacterial lawn where bacterial density was highest (Fig. 1C, Fig. 1-S1A-D). As a result, the tip of the animal’s head encountered the lawn boundary approximately once per minute. A few of these encounters resulted in a lawn leaving event, in which the animal exited the bacterial lawn (Fig. 1D, Fig. 1 – supplemental video 1). Lawn-leaving occurred in 26% of the assays, with a mean value of one event per 95 minutes (Fig. 1H-I, Supplementary Table 1).

**Figure 1:**
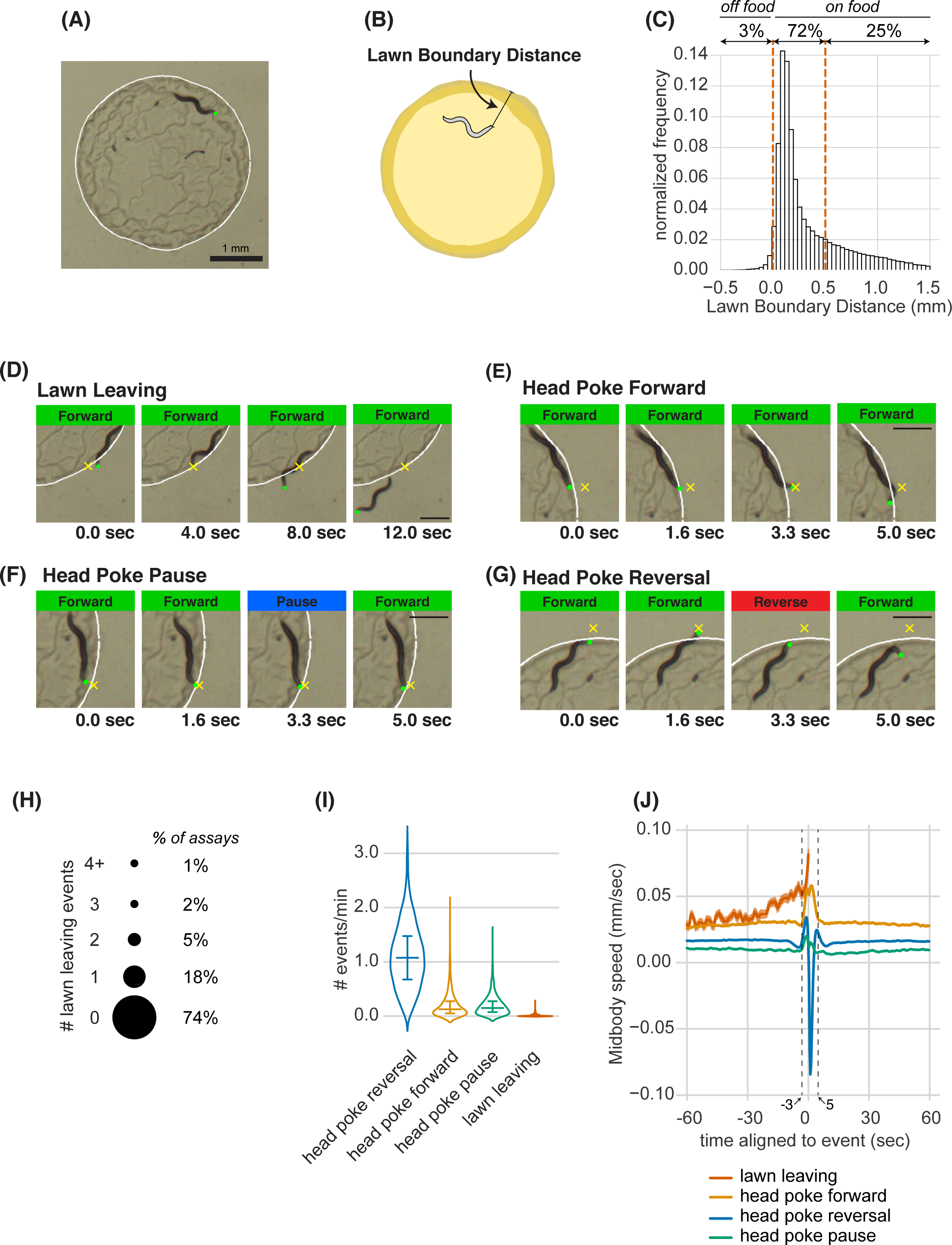
Animals make foraging decisions at the boundaries of bacterial food lawns. **(A)** Image of *C. elegans* on a small lawn of bacteria. Head is indicated with a green dot, lawn boundary is indicated with a white line. Scale bar is 1 mm. **(B)** Schematic depicting lawn boundary distance measured as the distance from the head to the closest point on the lawn boundary. **(C)** Empirical distribution of lawn boundary distance in wild type datasets. Positive values indicate distances inside the lawn, negative values indicate distances outside the lawn. **(D-G)** Four example images from behavioral sequences generating different types of foraging decisions. Scale bar is 0.5 mm in all panels. Head is indicated with a green dot in all panels. In E-G, yellow X indicates the maximum displacement of the head outside the lawn during head pokes. **(D)** Lawn leaving occurs when an animal approaches lawn boundary and fully crosses through it to explore the bacteria-free agar outside the lawn. Yellow X indicates the position on the lawn boundary encountered by worm’s head before exiting the lawn. **(E)** Head Poke Forward occurs when animal continues forward movement on the lawn during and after poking its head outside the lawn. **(F)** Head Poke Pause occurs when animal pauses following head poke before resuming forward movement on the lawn. **(G)** Head Poke Reversal is generated by a reversal following head poke before resuming forward movement on the lawn. **(H)** Number of lawn leaving events per animal in 40 minute assay. Bubble sizes are proportional to the percentage of animals that execute each number of lawn leaving events within a 40 minute assay. **(I)** Frequency of different foraging decisions. **(J)** Midbody speed aligned to different foraging decision types. Wild type dataset (C,H,I,J): n = 1586 animals. Violin plots in (I) show median and interquartile range. In all time-averages, dark line represents the mean and shaded region represents the standard error. See Supplementary Table 1.

Once an animal left the lawn, it typically remained outside of the food for 1-2 minutes before re-entering the lawn (Fig. 1-S1E, Supplementary Table 1). Most edge encounters were not followed by leaving; instead, animals that poked their head outside of the lawn either continued forward locomotion (head poke-forward), paused (head poke-pause), or executed a reversal (head poke-reversal) (Fig. 1E-G, Fig. 1 – supplemental video 1). Head poke-reversals were the most common event after lawn edge encounter (1.1 events/min), followed by head poke-forward and head poke-pause events; lawn leaving events were the least frequent (Fig. 1I, Supplementary Table 1).

High resolution analysis revealed characteristic behavioral dynamics prior to different events. Each head poke encounter with the lawn edge was preceded by an increase in speed over three seconds, and resolved by a reversal, pause, or continued forward movement within 5 seconds (Fig. 1J). Lawn leaving events encompassed a longer acceleration that began 30 seconds before the leaving event (Fig. 1J). At longer time scales, the average speed over one minute before and after the edge encounter varied slightly for different classes of events, with head poke-forward events and lawn-leaving associated with the highest speeds. The different behavioral patterns preceding lawn leaving and head pokes suggest that lawn leaving is a distinct foraging decision, and not a random resolution of an edge encounter. In particular, it suggests that lawn leaving is linked to a persistent behavioral state reflected in ongoing locomotion speed.

### Lawn leaving behavior is associated with high arousal states on short and long timescales

To gain further insight into the behavioral states preceding lawn leaving, we coarse-grained our behavioral measurements into 10 second intervals (“bins”) and expanded the time axis to allow analysis across different durations. On average, a slow rise in speed began two minutes before lawn leaving, with a rapid acceleration in the last minute before leaving (Fig. 2A). We analyzed these behaviors in the context of the well-characterized roaming and dwelling states (Ben Arous et al., 2009; Flavell et al., 2013; Fujiwara et al., 2002). Training a two-state Hidden Markov Model (HMM) based on absolute speed and angular speed (reorientations) recovered a bifurcation of roaming and dwelling states (Fig. 2-S1A-D) (Methods). Using these criteria, wild type animals on small lawns roamed for a median of 12% of the duration of the assay (Fig. 2-S1C, Supplementary Table 2). Although 80% of lawn leaving events occurred within roaming states, less than 1% of the 10 second roaming bins contained a lawn leaving event (Fig. 2B, Supplementary Table 2). Five minutes before leaving, 34% of animals were roaming, a fraction that steadily increased such that 80% of animals were roaming immediately before leaving (Fig. 2C, Supplementary Table 2). By contrast, the fraction of animals roaming before head poke-reversals increased only slightly before the event (Fig. 2-S3C). Within roaming states, the average speed was constant until the rapid acceleration ∼30 seconds prior to lawn leaving (Fig. 2D). To ask if the brief acceleration accounts for the apparent increase in roaming before lawn leaving, we compared the duration of roaming states that preceded lawn leaving to those that did not. We found that roaming states directly before lawn leaving were slightly longer than other roaming state, arguing that acceleration events alone do not drive the correlation between roaming and lawn leaving (Fig. 2-S1E).

**Figure 2:**
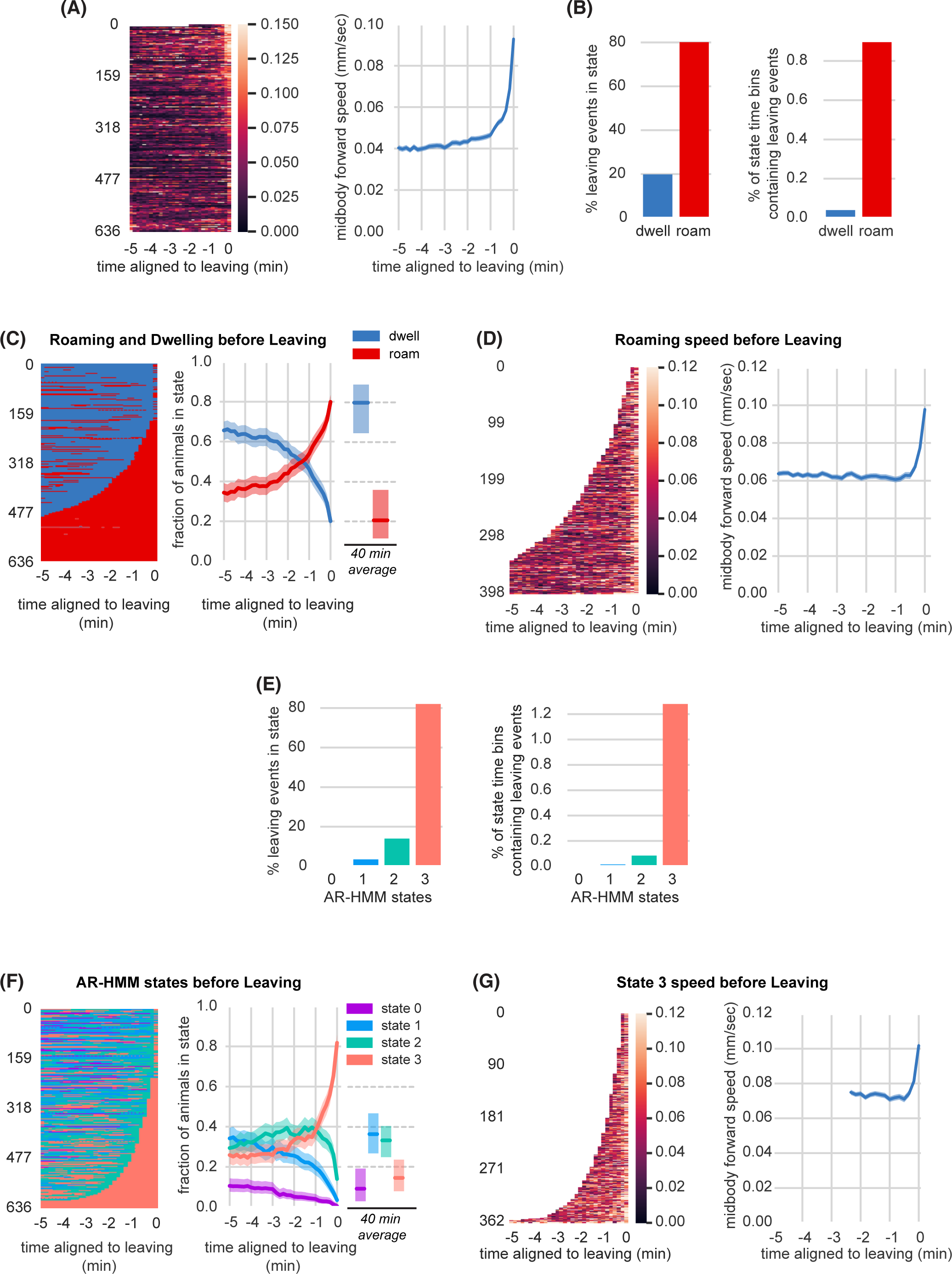
Lawn leaving is associated with high arousal states on multiple timescales. **(A)** Midbody forward speed aligned to lawn leaving. Left, heatmap of individual speed traces. Right, mean midbody forward speed computed across the heatmap traces. White space indicates missing data or times when animal was off the lawn. **(B)** Overlap of lawn leaving events with roaming and dwelling. Left, percentage of leaving events found in each state. Right, percentage of 10 s roaming or dwelling bins that contain a leaving event. **(C)** Roaming and dwelling aligned to lawn leaving. Left, heatmap of roaming/dwelling state classifications aligned to lawn leaving event. Center, fraction of animals in roaming or dwelling state prior to leaving. Right, total fraction of time spent roaming and dwelling in all assays that included a lawn-leaving event (n=371). Median is highlighted and interquartile range is indicated by shaded area. **(D)** Roaming speed aligned to lawn leaving. Left, heatmap showing speed of roaming animals before lawn leaving. Right, mean roaming speed computed at times when less than 10% of aligned traces had missing data. White space indicates times when animals were not roaming before leaving. **(E)** Overlap of lawn leaving events with AR-HMM states. Left, percentage of leaving events found in each state. Right, percentage of 10 s bins per AR-HMM state that contain a leaving event. **(F)** AR-HMM states aligned to lawn leaving. Left, heatmap of AR-HMM state classifications. Center, fraction of animals in each state prior to leaving. Right, total fraction of time spent in each AR-HMM state in all assays that included a lawn-leaving event (n=371). Median is highlighted and interquartile range is indicated by shaded area. **(G)** State 3 speed aligned to lawn leaving. Left, heatmap showing speed of animals in state 3 before lawn leaving. Right, mean state 3 speed computed at times when less than 10% of aligned traces had missing data. White space indicates times when animals were not in state 3 before leaving. See Supplementary Table 2.

The roaming-dwelling HMM was initially developed on uniform bacterial lawns (Flavell et al., 2013), which elicit more roaming than the small lawns used here (Fig. 2-S1F-H, Supplementary Table 2). Expanding this model to include posture information identified eight subclasses of dwelling states (Cermak et al., 2020). To explore alternative analysis methods, we generated an HMM behavioral state model trained on our experimental conditions. An Autoregressive Hidden Markov model (AR-HMM) (Buchanan et al., 2017; Linderman et al., 2020), trained over an expanded set of locomotion parameters, converged on four states that largely reflect forward locomotion speed and related features (Fig. 2-S2A-K, Methods). These four states provided an different segmentation of locomotory arousal that partly overlaps with roaming and dwelling states (Fig. 2-S2M-N). Most lawn leaving events occurred during the high speed, high arousal AR-HMM state 3 (Fig. 2E). Animals spent a median of 9% of the time in state 3 (Fig 2-S2K, Supplementary Table 2), but five minutes before leaving, 26% of animals were in state 3, which ramped to 82% immediately before leaving (Fig. 2F, Supplementary Table 2). Like roaming, state 3 increased only slightly before head poke reversals (Fig. 2-S3E). Animals across all behavioral states – roaming, dwelling, and the four AR-HMM states – were similarly distributed across the bacterial lawn, with strong enrichment near the lawn boundary (Fig. 2-S4A-D).

Both the two-state model and the four-state model show that arousal states increase before lawn leaving over at least two phases: an enrichment of a high arousal state 3-5 minutes before leaving, followed by a rapid acceleration in the last 30 seconds. Because the widely used two-state roaming-dwelling model captured the key features of lawn-leaving as well as these alternative methods, we applied that analysis to subsequent experiments.

### Food intake regulates arousal states and lawn leaving

Animals roam more and leave lawns more frequently when bacteria are hard to ingest or have low nutritional value (Ben Arous et al., 2009; Shtonda, 2006). Conversely, simply increasing food concentration suppressed roaming and leaving on small lawns (Fig. 3A-B, Supplementary Table 3) (Bendesky et al., 2011). To separate the behavioral effects deriving from sensation of food odors from ingestion of food, we treated *E. coli* OP50 bacteria with aztreonam, an antibiotic that renders bacteria inedible to *C. elegans* by preventing bacterial cell septation (Ben Arous et al., 2009; Gruninger et al., 2008). Both lawn leaving and roaming were dramatically increased on aztreonam-treated *E. coli* lawns (Fig. 3C-D, Fig. 3-S1A-B, Supplementary Table 3, Table 2). Animals on aztreonam-treated lawns maintained the characteristic speed acceleration in the last 30 seconds before leaving (Fig. 3-S1C).

**Figure 3:**
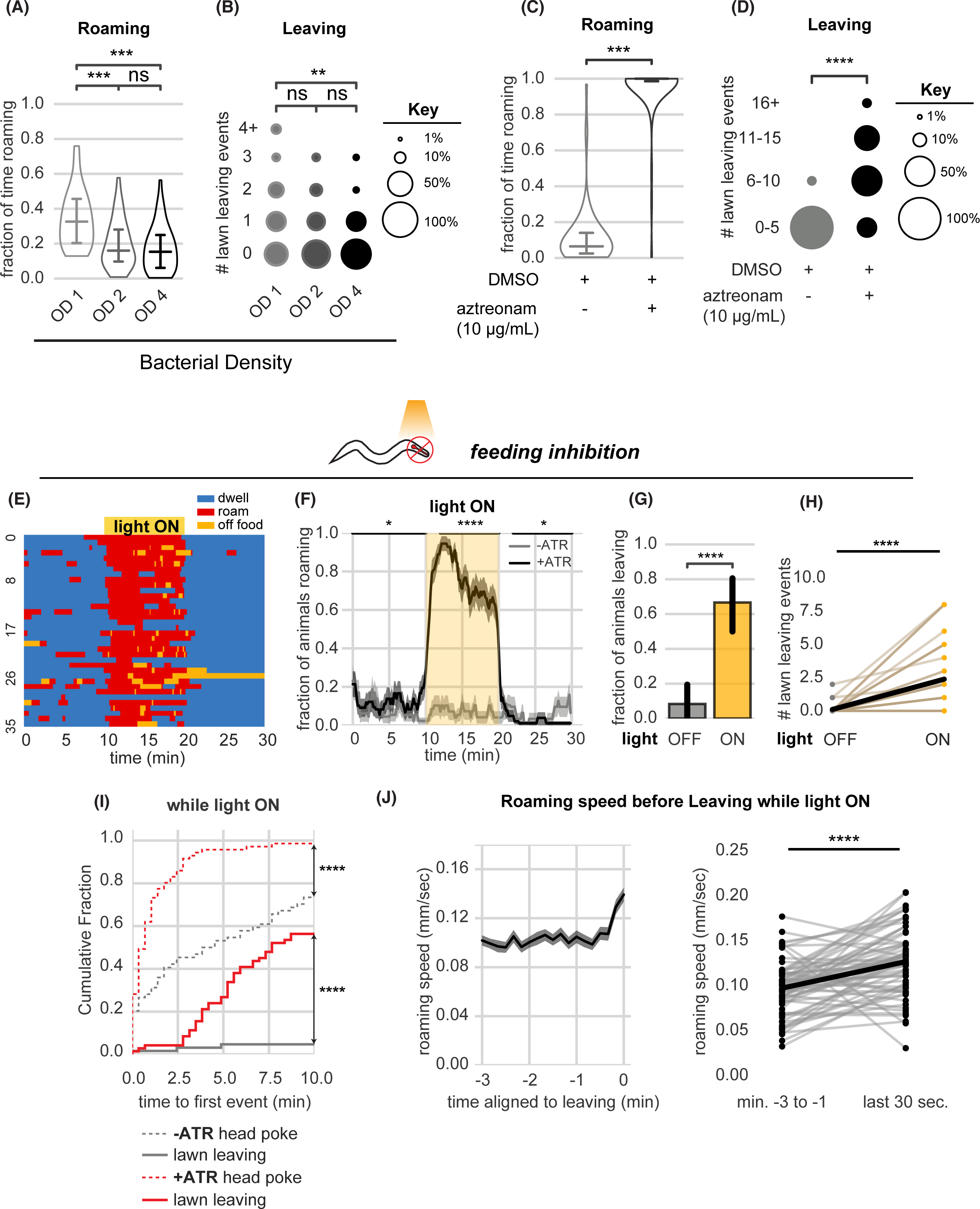
Food intake regulates arousal states and lawn leaving. **(A-B)** Increasing bacterial density suppresses roaming and leaving. **(A)** Fraction of time roaming on small lawns seeded with bacteria of different optical density (OD). OD1: n = 45, OD2: n = 46, OD4: n = 47, Statistics by one-way ANOVA with Tukey’s post-hoc test on logit-transformed data. **(B)** Number of lawn leaving events per animal in the same assays as (A). Statistics by Kruskal-Wallis test with Dunn’s multiple comparisons test. **(C-D)** Animals on inedible food roam and leave lawns more than animals on edible food. **(C)** Fraction of time roaming on inedible bacteria generated by adding aztreonam dissolved in DMSO to *E. coli* growing on plates. +DMSO/+aztreonam n = 64, +DMSO (control) n = 66, Statistics by Student’s t-test performed on logit-transformed data. **(D)** Number of lawn leaving events in the same assays as (C). Statistics by Mann-Whitney U-test. **(E-J)** Feeding inhibition by optogenetic depolarization of pharyngeal muscles stimulates roaming and lawn leaving. **(E)** Heatmap showing roaming and dwelling for animals before, during, and after 10-minute optogenetic feeding inhibition. Data for animals pre-treated with all-trans retinal (+ATR) is shown. **(F)** Fraction of animals roaming before, during, and after optogenetic feeding inhibition. Light ON period denoted by yellow shading (+ATR n = 36). Control animals not pre-treated with all-trans retinal (-ATR n = 32). Statistics by Student’s t-test comparing +/-ATR data averaged and logit-transformed during intervals indicated by black lines above plots: Minutes 0-10, 12-20, 22-30. **(G)** A greater fraction of animals leave lawns during feeding inhibition. Statistics by Fisher’s exact test. **(H)** Number of lawn leaving events in the same assays as (G). Statistics by Wilcoxon rank-sum test. **(I)** Cumulative distribution of time until the first head poke reversal or lawn leaving event during feeding inhibition. Statistics by Kolmogorov-Smirnov 2-sample test. **(J)** Roaming animals accelerate before leaving during feeding inhibition. Left, mean roaming speed of animals before leaving. Right, quantification of roaming speed increase from minutes −3 to −1 to the last 30 second before leaving. Statistics by Wilcoxon rank-sum test. Statistics: ns, not significant, * p < 0.05, *** p < 10_-3_,**** p < 10^-4^ In time-averages (F,J), dark line represents the mean and shaded region represents the standard error. Violin plots show median and interquartile range. In (H), each dot pair connected by a line represents data from a single animal. In (J), each dot pair connected by a line represents data preceding a single lawn leaving event. Thick black line indicates the average. See Supplementary Table 3.

Locomotion speed is regulated by direct interoception of pharyngeal pumping, the motor behavior associated with intake of bacterial food (Rhoades et al., 2019). To ascertain whether similar mechanisms affect arousal and lawn leaving, we acutely inhibited pharyngeal pumping by optogenetic activation of the red-shifted channelrhodopsin ReaChR in pharyngeal muscle (Lin et al., 2013) (Fig. 3E-J). Acute feeding inhibition strongly stimulated both roaming and lawn leaving (Fig. 3E-H, Fig. 3-S1D-E, Supplementary Table 3, Table 2).

While roaming and head pokes began immediately upon feeding inhibition, leaving events did not occur immediately but instead accumulated throughout the 10-minute stimulation interval (Fig. 3I). In each case, leaving was preceded by a 30 second acceleration in speed (Fig. 3J). Thus, feeding inhibition elicits both roaming and lawn-leaving behaviors, and lawn-leaving occur probabilistically during roaming states.

### Parallel regulation of arousal states and leaving by neuromodulatory mutants

To further examine the relationship between arousal states and lawn leaving, we examined mutants with known alterations in roaming and dwelling. Animals deficient in serotonin *(tph-1)*, dopamine *(cat-2)*, or the neuropeptide receptor NPR-1 *(npr-1)* roam at a higher rate than wild type (Cermak et al., 2020; Cheung et al., 2005; Flavell et al., 2013; Omura et al., 2012; Sawin et al., 2000). We found that each of these mutants showed increased lawn leaving compared to wild-type controls, strengthening the observed correlation between roaming and leaving rates (Fig 4A-C, E-G, Table 2). In addition, the fraction of animals roaming increased over several minutes before leaving events, as was observed in wild type animals (Fig. 4I-K). Finally, each mutant accelerated during the 30 seconds prior to leaving (Fig. 4M-O). Thus, although the molecular basis of arousal is different in each of these mutants, the overall dynamics of roaming and lawn leaving are preserved across genotypes.

**Figure 4:**
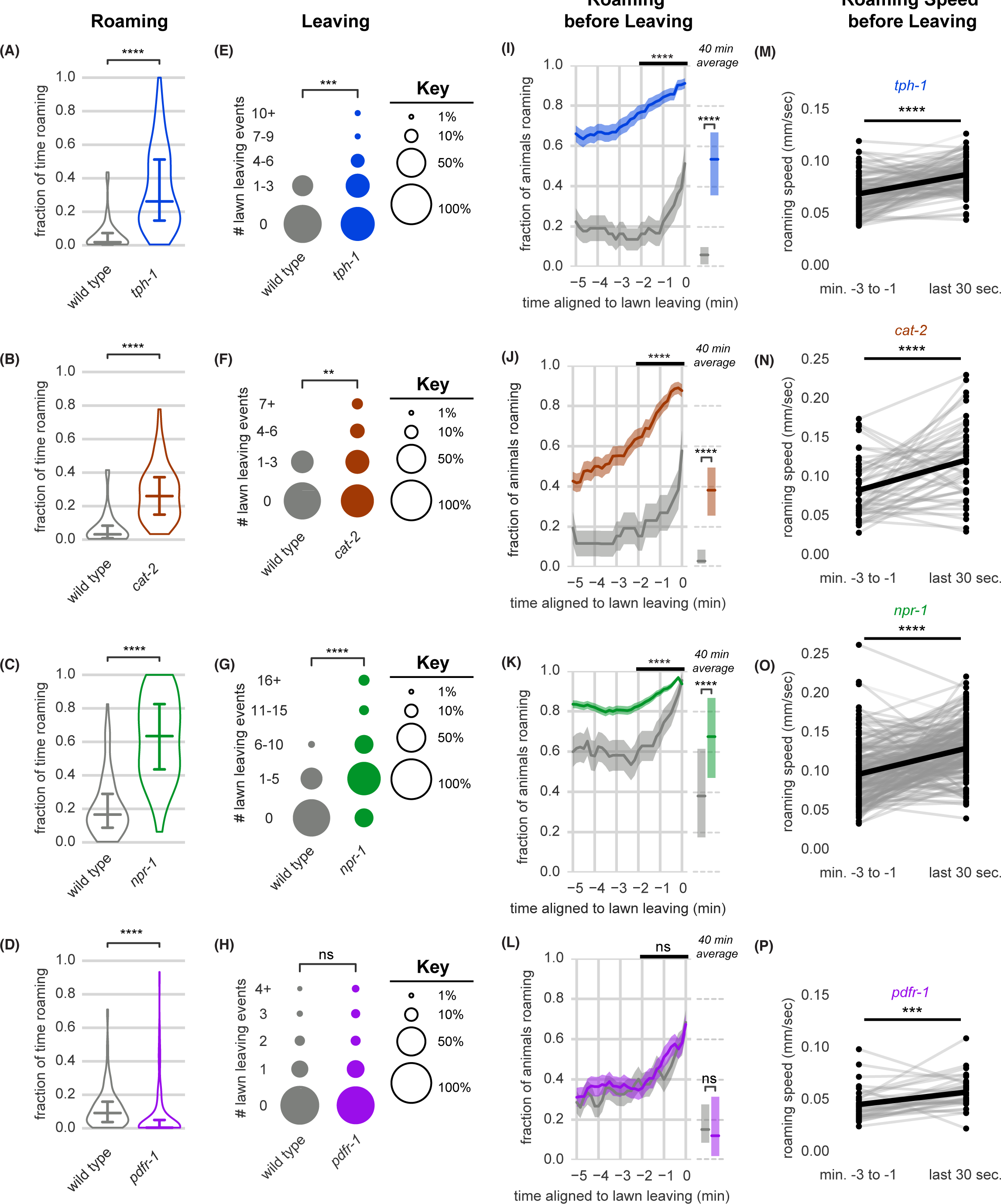
Neuromodulatory signaling mutants retain coupling of arousal and leaving. **(A-P)** Roaming, lawn leaving, roaming before leaving, and roaming speed quantified across mutants in four neuromodulatory genes known to alter roaming and dwelling: *tph-1*, *cat-2*, *npr-1*, and *pdfr-1*. **(A-D)** Fraction of time roaming. Statistics by Student’s t-test on logit-transformed data. **(E-H)** Number of lawn leaving events per animal. Statistics by Mann-Whitney U test. **(I-L)** Fraction of animals roaming before lawn leaving. Left, fraction of animals roaming in the last 5 minutes before lawn leaving. WT and mutant roaming fractions were compared over the two minutes prior to leaving (black bar). Right, total fraction of time spent roaming and dwelling in all assays that included a lawn-leaving event. Statistics by Student’s t-test on logit-transformed data. Note that roaming levels were unusually low (A,B,I,J) or high (C,K) in the wild-type controls for these groups, and therefore genotypes cannot be compared across different experimental panels. **(M-P)** Roaming speed before leaving computed at times when less than 10% of aligned traces had missing data. Paired plots indicate the average speed from minutes −3 to −1 and in the last 30 seconds before leaving per animal. Statistics by Wilcoxon rank-sum test. Statistics: ns not significant (p > 0.05), ** p < 0.01, *** p < 10^-3^, **** p < 10^-4^ Violin plots and box plots show median and interquartile range. In time-averages, dark line represents the mean and shaded region represents the standard error. In paired plots (M-P), each dot pair connected by a line represents data preceding a single lawn leaving event. Thick black line indicates the average. (*tph-1* n = 139, wild type controls n = 127; *cat-2* n = 88, wild type controls n = 76; *npr-1* n = 91, wild type controls n = 90; *pdfr-1* n = 265, wild type controls n = 247). See Supplementary Table 4.

**Table 2a:**
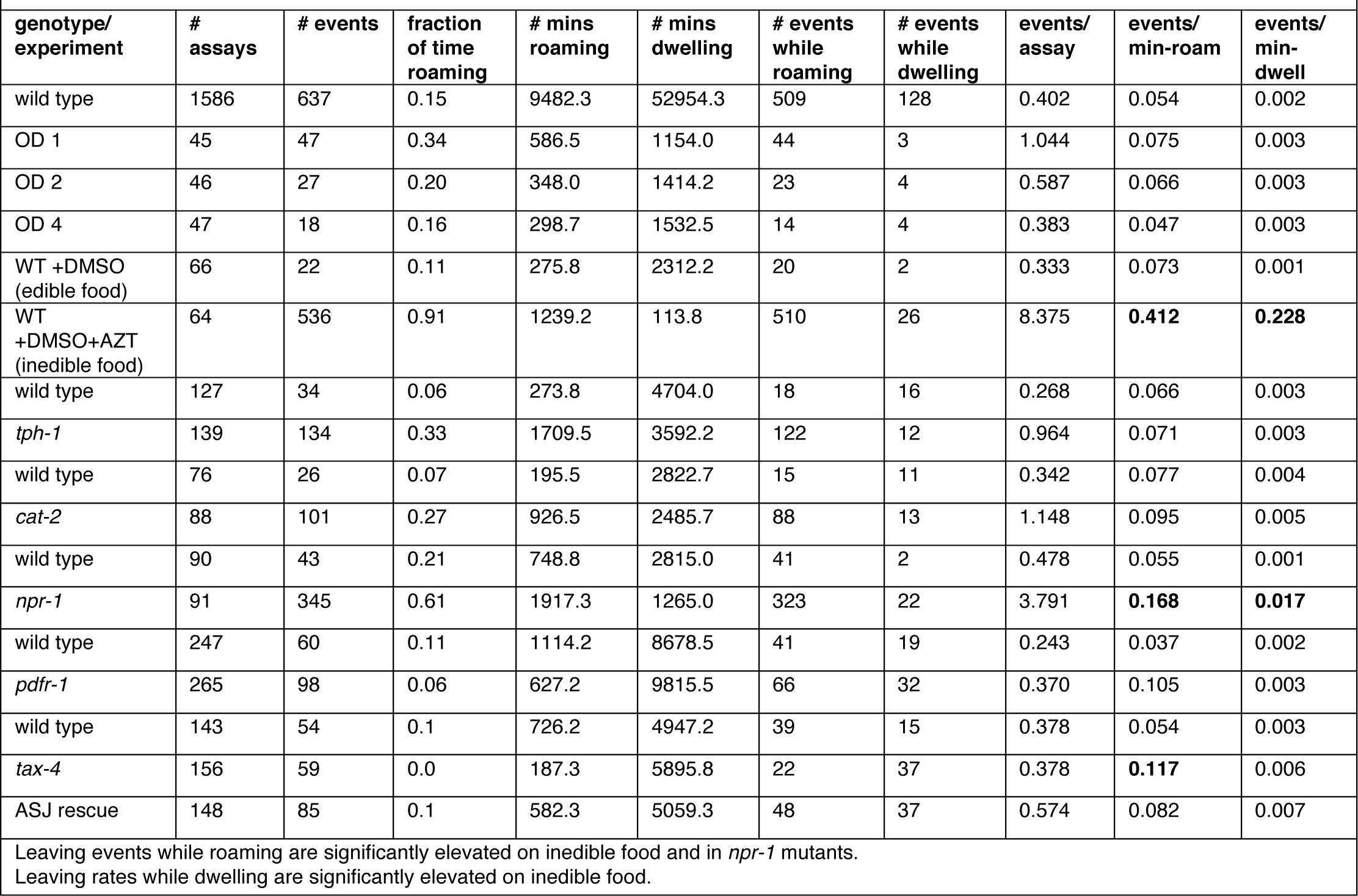
Lawn Leaving Events Per minute Roaming/Dwelling.

**Table 2b:**
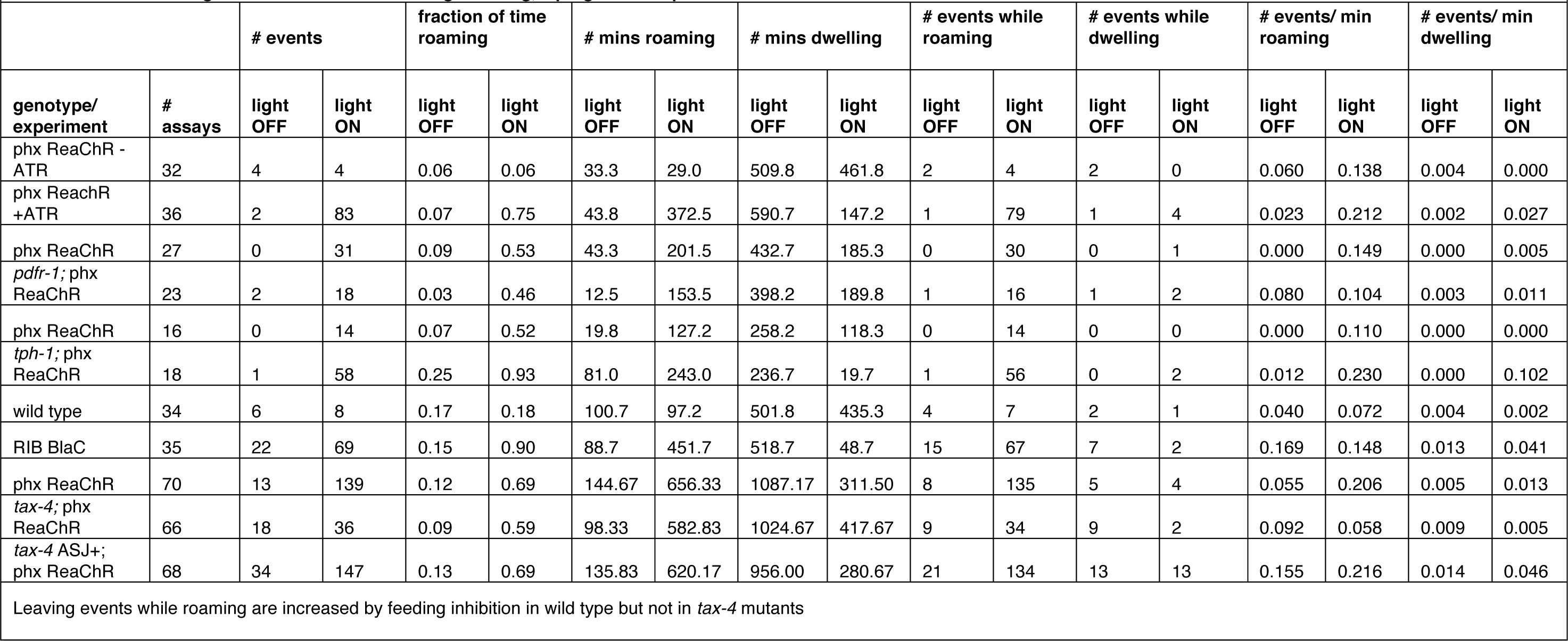
Lawn Leaving Events Per minute Roaming/Dwelling, Optogenetic Experiments.

Animals lacking the G-protein coupled receptor Pigment Dispersing Factor Receptor (PDFR-1) roam less than wild type (Flavell et al., 2013; Ji et al., 2021; Meelkop et al., 2012). On the small lawns used here, *pdfr-1* mutants roamed less, but left food lawns at the same low rate as wild type (Fig. 4D,H). Roaming rates increased similarly during the 3 minutes prior to lawn leaving in wild type and *pdfr-1* animals, suggesting that the coupling of roaming and leaving does not require PDFR-1 signaling (Fig. 4L). Although their basal locomotion speed is lower (Flavell et al., 2013; Ji et al., 2021), *pdfr-1* did accelerate slightly before lawn leaving (Fig. 4P). In summary, neuromodulatory mutants varied in the fraction of time spent roaming and dwelling, but in each case lawn-leaving behaviors were coupled to roaming and a brief speed acceleration.

Optogenetic inhibition of feeding elicited immediate roaming and probabilistic lawn leaving in both *pdfr-1* and *tph-1* mutants (Fig. 4-S1), indicating that neither of these neuromodulators is essential for interpreting feeding inhibition. This was surprising because the *tph-1-*expressing NSM neurons sense bacterial ingestion and signal food availability via serotonin release (Rhoades et al., 2019), and were therefore candidates to relay feeding signals. NSM may act through additional transmitters as well as serotonin, or it may be redundant with additional neurons that detect feeding inhibition.

### Acute circuit manipulation drives deterministic roaming and probabilistic leaving

As a complement to the neuromodulatory mutants, we employed a circuit-based approach to manipulate arousal levels and examine effects on lawn leaving. Roaming is strongly stimulated by *pdfr-1;* we defined sites of *pdfr-1* expression that stimulate roaming using an intersectional Cre-Lox system that restores *pdfr-1* expression in targeted groups of cells in *pdfr-1* mutant animals (Fig. 5-S1, Fig. 5-S2A) (Flavell et al., 2013). Previous work identified a moderate effect of AIY, RIM, and RIA neurons as mediators of *pdfr-1-*dependent roaming (Flavell et al., 2013). We observed a stronger rescue of roaming upon *pdfr-1* expression solely in the RIB neurons, which are active during rapid forward locomotion (Fig. 5-S2B-D) (Ji et al., 2021; Wang et al., 2020).

Following this result, we asked whether optogenetic activation of RIB might be sufficient for roaming in wild type animals. PDFR-1 signals through the heterotrimeric G protein Gαs to increase cAMP levels (Janssen et al., 2008), and optogenetic activation of groups of PDFR-1-expressing neurons with the bacterial light-activated adenylyl cyclase BlaC increases roaming (Flavell et al., 2013; Ryu et al., 2010). We found that acute optogenetic activation of only RIB with BlaC induced immediate roaming in over 80% of animals (Fig. 5A-C, Fig. 5-S3A-B).

**Figure 5:**
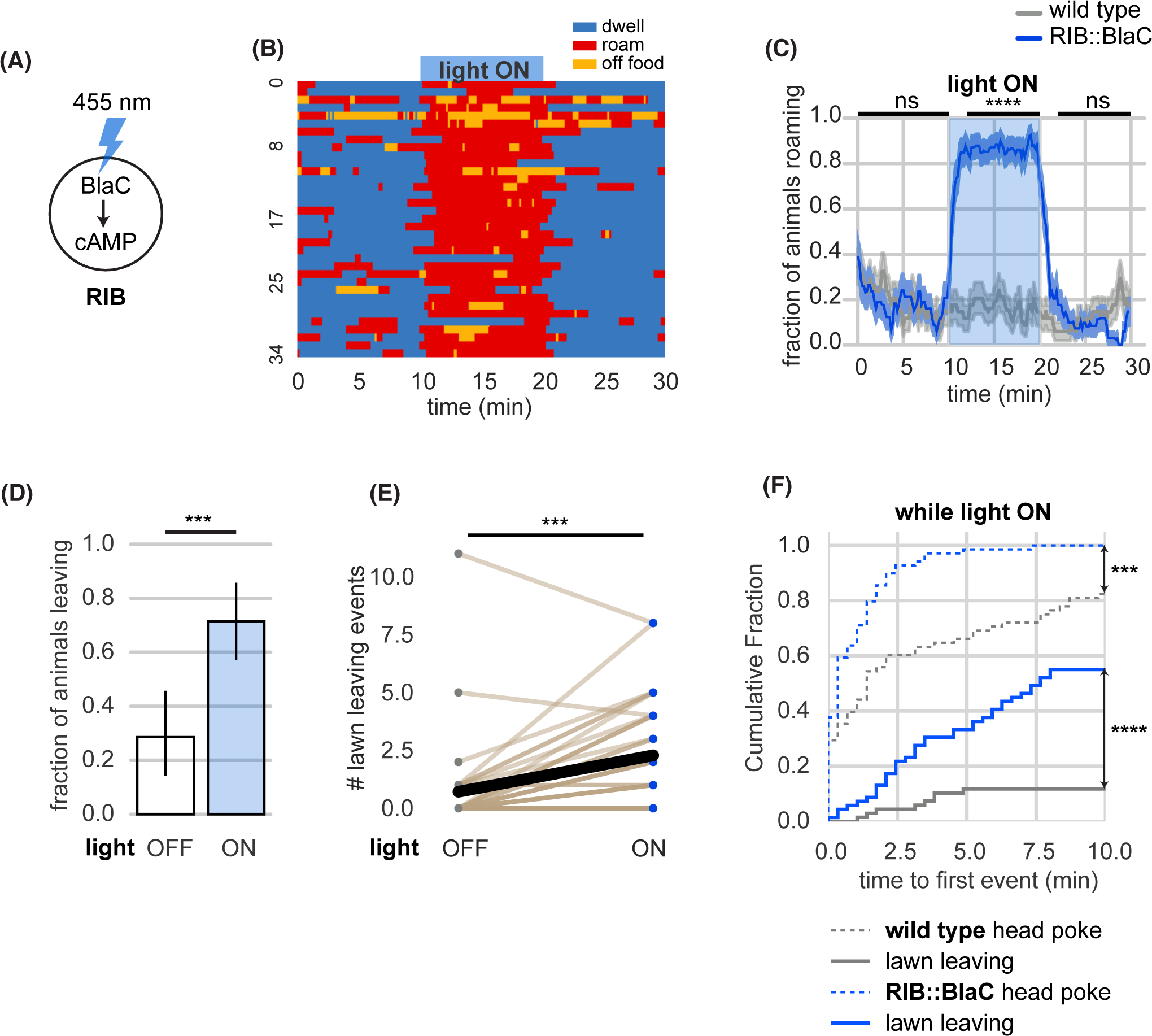
Acute circuit manipulation drives deterministic roaming and probabilistic leaving. **(A)** Experimental design. Stimulation of bacterial light-activated adenylyl cyclase (BlaC) with 455 nm light increases cAMP synthesis in RIB::BlaC neurons. **(B)** Heatmap showing roaming and dwelling for RIB::BlaC animals before, during, and after 10-minute optogenetic stimulation. **(C)** Fraction of animals roaming before, during, and after optogenetic blue light stimulation. Light ON period denoted by blue shading. Control animals are wild type. Statistics by Student’s t-test comparing RIB::BlaC and wild type animals averaged and logit-transformed during intervals indicated by black lines above plots: Data compared at 0-10, 12-20, 22-30 minutes. **(D)** A greater fraction of RIB::BlaC animals leave lawns when the light is ON vs. OFF. Statistics by Fisher’s exact test. **(E)** Number of lawn leaving events under the same conditions as (D). Statistics by Wilcoxon rank-sum test. **(F)** Cumulative distribution of time until the first head poke reversal or lawn leaving event while the light is ON. Statistics by Kolmogorov-Smirnov 2-sample test. Statistics: ns not significant (p > 0.05), ** p < 0.01, *** p < 10^-3^, **** p < 10^-4^ In time-averages (C), dark line represents the mean and shaded region represents the standard error. (RIB::BlaC n = 35, wild type n = 34) In (E), each dot pair connected by a line represents data from a single animal. Thick black line indicates the average. See Supplementary Table 5.

RIB::BlaC stimulation also potentiated lawn leaving, with over 70% of animals leaving lawns during the stimulation period (Fig. 5D-E). Like lawn-leaving during optogenetic feeding inhibition, this behavior was probabilistic across the 10-minute light pulse (Fig. 5F). Animals accelerated slightly in the 30 seconds before leaving, but the speed increase appeared less pronounced than in other manipulations, suggesting that RIB activity may partially occlude the acceleration motif in lawn leaving (Fig. 5-S3F). Neither roaming nor lawn leaving was potentiated by blue light exposure in wild type control animals not expressing BlaC (Fig. 5C, Fig. 5-S3A, C-D).

### Chemosensory neurons couple roaming dynamics, internal state, and lawn leaving

In addition to the neuromodulatory arousal systems, multiple food- and pheromone-sensing chemosensory neurons affect roaming, dwelling, and leaving behaviors (Table 1). Most of these neurons use the cyclic nucleotide-gated channel gene *tax-4* for sensory transduction, and *tax-4* mutants have diminished roaming behavior (Ben Arous et al., 2009; Fujiwara et al., 2002) (Fig. 6A, Table 1). However, we found that *tax-4* mutant animals continued to leave lawns – indeed, they left at slightly higher rates than wild-type animals (Fig. 6B, Table 2). Moreover, the temporal relationship between roaming and leaving was altered in *tax-4* mutants, which typically roamed for only ∼1 minute prior to lawn leaving (Fig. 6C, Fig. 6-S1A-B). These results indicate that loss of *tax-4* disrupted the characteristic arousal dynamics associated with leaving.

**Figure 6:**
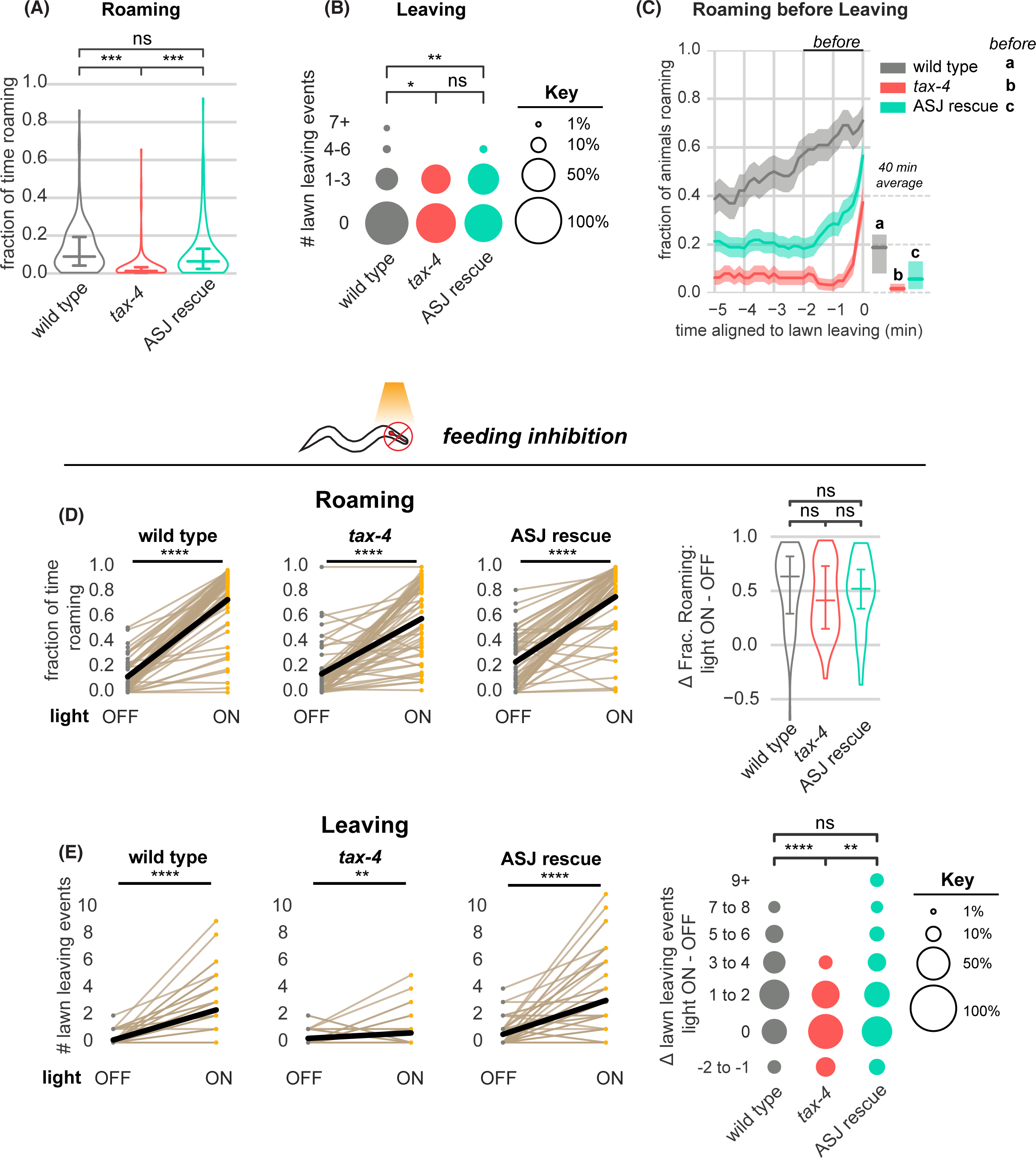
*tax-4*-expressing sensory neurons couple roaming and lawn leaving. **(A-C)** Roaming and leaving in *tax-4* mutants, and rescue by *tax-4* expression in ASJ neurons (wild type n = 143, *tax-4* n = 156, *tax-4* ASJ rescue n = 148). Additional features of roaming and leaving are shown in Fig. 6-S1; results with additional rescued neurons are in Fig. 6-S2. **(A)** Fraction of time roaming. Statistical tests, one-way ANOVA followed by Tukey’s post hoc test on logit-transformed data. **(B)** Number of lawn-leaving events per animal. Statistical tests, Kruskal-Wallis followed by Dunn’s multiple comparisons test. **(C)** Fraction of animals roaming before lawn-leaving. Left, fraction of animals roaming in the last 5 minutes before lawn leaving. Roaming fractions were compared over the two minutes prior to leaving (black bar). Right, total fraction of time spent roaming and dwelling in all assays that included a lawn-leaving event. Key, statistical tests of differences in roaming two minutes before leaving by one-way ANOVA followed by Tukey’s post hoc test on logit-transformed data. **(D-E)** Optogenetic feeding inhibition in *tax-4* mutants and rescued strains (wild type n = 70, *tax-4* n = 66, *tax-4* ASJ rescue n = 68). **(D)** Left, Fraction of time spent roaming under optogenetic feeding inhibition in *tax-4* and ASJ rescue strain. Statistics by paired t-test on logit-transformed data. Right, Difference in the fraction of time roaming when the light is ON and OFF. Statistics by one-way ANOVA followed by Tukey’s post hoc test on logit-transformed data. **(E)** Left, Number of lawn leaving events under optogenetic feeding inhibition in *tax-4* and ASJ rescue strain. Statistics by Wilcoxon rank-sum test. Right, Difference in the number of lawn leaving events when the light is ON and OFF. Statistics by Kruskal-Wallis test followed by Dunn’s multiple comparisons test. ns not significant (p > 0.05), * p < 0.05, *** p < 10^-3^, **** p < 10^-4^ Violin plots and box plots (A,D) show median and interquartile range. In time-averages (C), dark line represents the mean and shaded region represents the standard error. In paired plots (D,E), each dot pair connected by a line represents data from a single animal. Thick black line indicates the average. See Supplementary Table 6.

Many of the 15 classes of sensory neurons that express *tax-4* have been implicated in roaming or leaving behaviors (Table 1). We rescued *tax-4* separately in AWC, which senses food odors; ASK, which senses amino acids and pheromones; ASJ and ASI, which sense pheromones, food, and toxins; and URX/AQR/PQR, which sense environmental oxygen (Ben Arous et al., 2009; Bendesky et al., 2011; Greene et al., 2016; Milward et al., 2011). Significant effects on roaming or leaving behavior were observed upon ASJ, ASK, or AWC rescue (Fig. 6A-C, Fig. 6-S1 and S2). The strongest effects resulted from *tax-4* rescue in the ASJ neurons, which partially restored roaming levels before lawn leaving (Fig. 6A-C, Fig 6-S1), and *tax-4* rescue in the ASK neurons, which paradoxically suppressed leaving to a level below that of either wild type or *tax-4* animals (Fig 6-S2). Because ASJ and ASK showed opposite effects in these experiments, we also examined strains in which both neurons were rescued. Combined ASJ and ASK rescue normalized roaming and leaving compared to ASK rescue alone, albeit not to fully wild-type levels (Fig 6-S2). While we have not tested all neurons and combinations, these results suggest that multiple *tax-4-*expressing sensory neurons have roles in arousal-related behaviors and highlight ASJ as a regulator of roaming and leaving dynamics.

Next, we asked how *tax-4* mutants responded to acute inhibition of feeding. As in wild-type animals, optogenetic inhibition of pharyngeal pumping resulted in an immediate and strong increase in roaming in *tax-4* mutants (Fig. 6D, Fig. 6-S3A). However, feeding inhibition in *tax-4* mutants increased leaving only slightly, unlike in the wild-type (Figure 6E, Table 2). Both of these effects were rescued by expressing *tax-4* in ASJ neurons (Fig. 6C-E, Fig. 6-S3B-D). Similarly, *tax-4* animals on inedible food roamed at the same high rate as wild type animals but produced significantly fewer lawn leaving events (Fig 6-S4); rescuing *tax-4* in ASJ restored lawn leaving to wild-type levels. Thus *tax-4* sensory mutants uncouple leaving behavior from its normal context in multiple ways: they can leave edible food lawns without an extended roaming state, and they are less likely to leave when feeding is inhibited, even while they are roaming.

## DISCUSSION

Both in the wild and in laboratory settings, foraging locomotion patterns exhibit scale invariance, meaning that similar statistics of movement displacement and duration arise on short and long timescales (Ayala-Orozco et al., 2004; Proekt et al., 2012). Long term arousal changes are regulated by neuromodulators that signal widely throughout the brain (Flavell et al., 2022; Taghert & Nitabach, 2012; Weissbourd et al., 2014), whereas moment-by-moment decision-making involves fast sensorimotor circuits (Gold & Shadlen, 2007). Here we show that *C. elegans* couples food leaving decisions, which unfold over seconds, to high arousal roaming states that last minutes. Although food intake and neuromodulatory signaling both alter the frequency of sustained *C. elegans* roaming states, these changes do not disrupt the coupling of roaming to food leaving decisions. Instead, sensory neurons link roaming and leaving behaviors, integrating these behavioral motifs, their dynamic properties, and their regulation by food intake.

### Behavioral arousal alters probabilistic decision-making

From an ethological perspective, lawn-leaving is a classical foraging decision based on an animal’s assessment of the quality of a food patch (Charnov, 1976). Previous studies of lawn-leaving have identified features of the environment, like food availability, population density, and toxic repellents, that affect leaving probability as predicted by foraging theory. Here, we have extended these observations using the framework of computational neuroethology (Datta et al., 2019), examining behavior in detail across time to identify rare leaving events and the behavioral context in which they occurred. We found that leaving was tightly coupled to roaming both during spontaneous behaviors and after acute manipulations that induced roaming states. In each case, and in neuromodulatory mutants that altered roaming frequency, leaving occurred when animals were roaming, occurred probabilistically over minutes during the roaming state, and was preceded by a brief 30-second acceleration in speed. The stereotyped features of leaving behavior, which were not shared by other lawn edge encounters, suggest that it is a discrete behavioral motif associated with roaming states. Leaving can be viewed as a form of decision-making—a discrete action that drives an adaptive behavioral choice between alternatives, influenced by internal states such as arousal (Kennedy et al., 2014).

Roaming and dwelling comprise a widely-used framework for defining arousal states in *C. elegans,* although changing experimental conditions or analysis methods can reveal sub-states or alternative classification frameworks (this work and Cermak et al., 2020; Gallagher et al., 2013; Raizen et al., 2008; Van Buskirk & Sternberg, 2007). Classical ethology presents such internal states as hierarchical and mutually exclusive (Tinbergen, 1951), while molecular and circuit analysis has extended and refined these ideas (Asahina et al., 2014; Flavell et al., 2013; Hindmarsh Sten et al., 2021; Hong et al., 2014; Moy et al., 2015; Pan et al., 2012). Our results indicate that lawn leaving behaviors are coupled to the roaming state, but within this state they are relatively rare and apparently probabilistic. More detailed studies may reveal additional regulators, such as an individual’s sensory experience, that shape the pattern of leaving behavior (Coen et al., 2014).

### Sensory neurons integrate internal state and external sensation to guide foraging decisions

Our results identify two distinct biological mechanisms that regulate lawn leaving. First, leaving is coupled to arousal state. Leaving rates are 20-fold higher in roaming animals than in dwelling animals, and the leaving rates of most arousal mutants are largely explained by the fact that they spend more time roaming (Table 2). The second mechanism is controlled by *tax-4* sensory neurons, which shape the behavioral dynamics of roaming and leaving across a range of conditions and stimulate leaving during feeding inhibition.

At a straightforward level, sensory neurons are well-placed to evaluate environmental quality in the framework of foraging theory. For example, ASK and AWC neurons sense amino acids and food odors (Bargmann, 2006), while ASJ neurons sense pheromones and toxins (Greene et al., 2016; Meisel et al., 2014), and other sensory neurons detect distinct environmental features (Bargmann, 2006). Accordingly, many sensory neurons affect roaming, dwelling, or leaving (Table 1), and their relative importance varies with context, such as the presence of chemical repellents or pheromones that signal population density (Dal Bello et al., 2021; Greene et al., 2016; Milward et al., 2011; Pradel et al., 2007). Under the conditions used here, ASK neurons suppressed and ASJ neurons promoted roaming and leaving, respectively. Rescuing *tax-4* in ASJ partially restored features of normal leaving dynamics, including roaming before leaving.

Others have shown that chronic feeding deprivation across hours drives lawn leaving through the action of *tax-4* in several sensory neurons (Olofsson, 2014). An unexpected result obtained here was that *tax-4* sensory neurons were required to drive the high leaving rates after acute inhibition of feeding. Together, these results suggest that the sensory neurons are sites at which internal feeding cues, behavioral states, and specific foraging decisions are integrated. Additional experiments are needed to define the full set of sensory neurons that couple feeding inhibition to leaving, the nature of the interoceptive signal from pharyngeal pumping, and the readout of the sensory neurons. Pharyngeal interoception is mediated in part by the serotonergic NSM neurons (Rhoades et al., 2019), but we found that serotonin was not essential for the effects of acute feeding inhibition. With respect to interoception, some of the *tax-4-*expressing sensory neurons detect tyramine, one of several biogenic amines that regulate lawn leaving (Bendesky et al., 2011; O’Donnell et al., 2020), and many detect neuropeptides as well (Hapiak et al., 2013).

Sensory neurons in *C. elegans* and other animals are most often studied in the context of rapid behavioral responses, but they also have critical roles in endocrine signaling. For example, intrinsically photosensitive retinal ganglion cells drive circadian entrainment in mammals (Hattar et al., 2002) and in cichlid fish, visual stimuli from females are sufficient to induce an endocrine androgen response in males (O’Connell et al., 2013). We speculate that the combination of endocrine signals, neuropeptides, and fast transmitters released by sensory neurons couples roaming states to leaving states. The insulin peptides and DAF-7-TGF-beta protein that regulate roaming, dwelling, and leaving are primarily produced by *tax-4-* expressing sensory neurons, so these neurons bridge sensory and endocrine regulation of foraging (Taylor et al., 2021). Expression of these genes is regulated by pheromones and metabolic conditions, providing an additional layer beyond neural activity for long-term regulation of behavioral state. Many *tax-4* sensory neurons also release classical neurotransmitters such as glutamate (ASK, AWC) and acetylcholine (ASJ), and therefore have the potential to modulate rapid behaviors such as the lawn-leaving motif (Taylor et al., 2021). Future studies can test this hypothesis while determining the mechanisms by which sensory neurons integrate signals to trigger lawn leaving.

## METHODS

### Key Resources Table

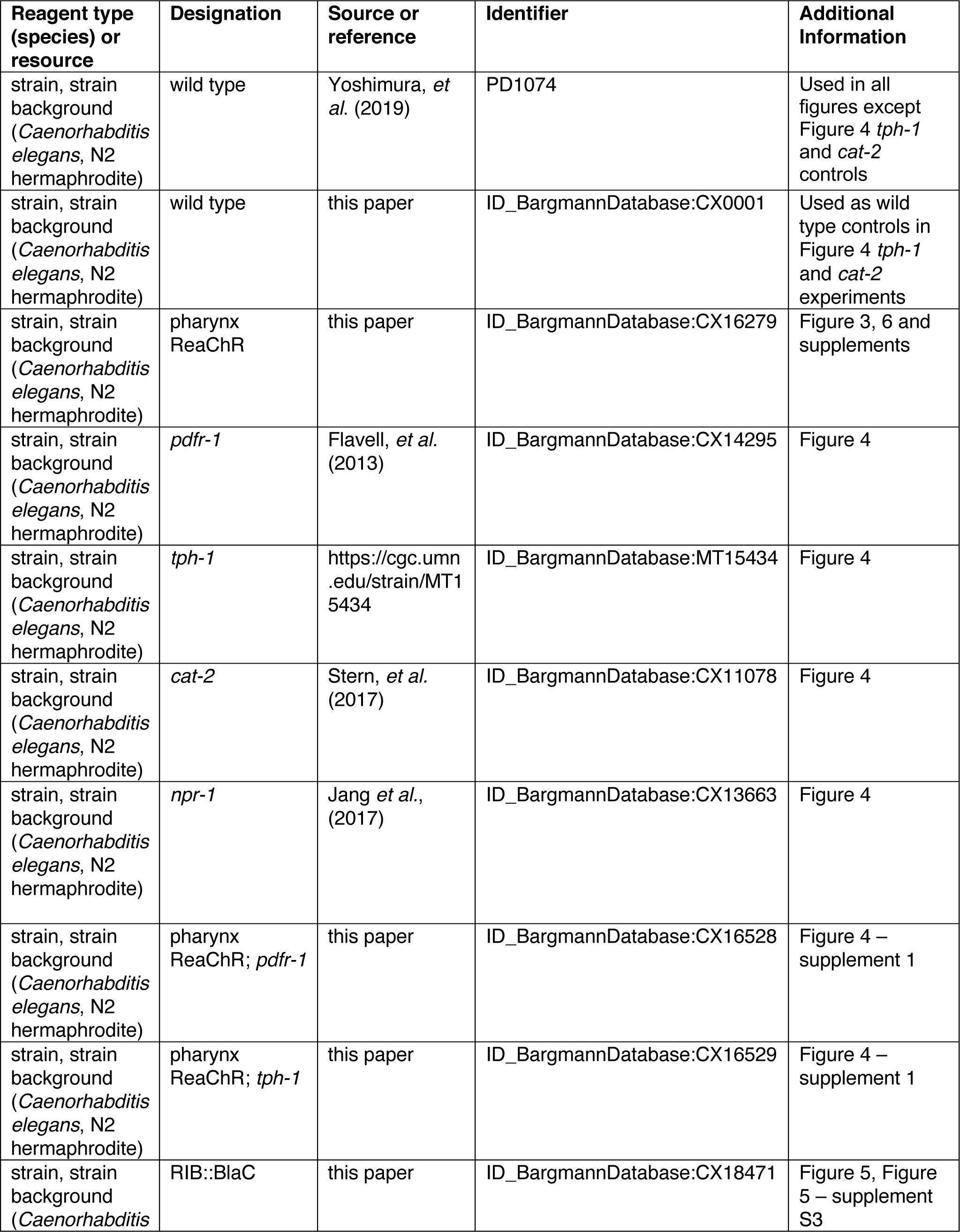
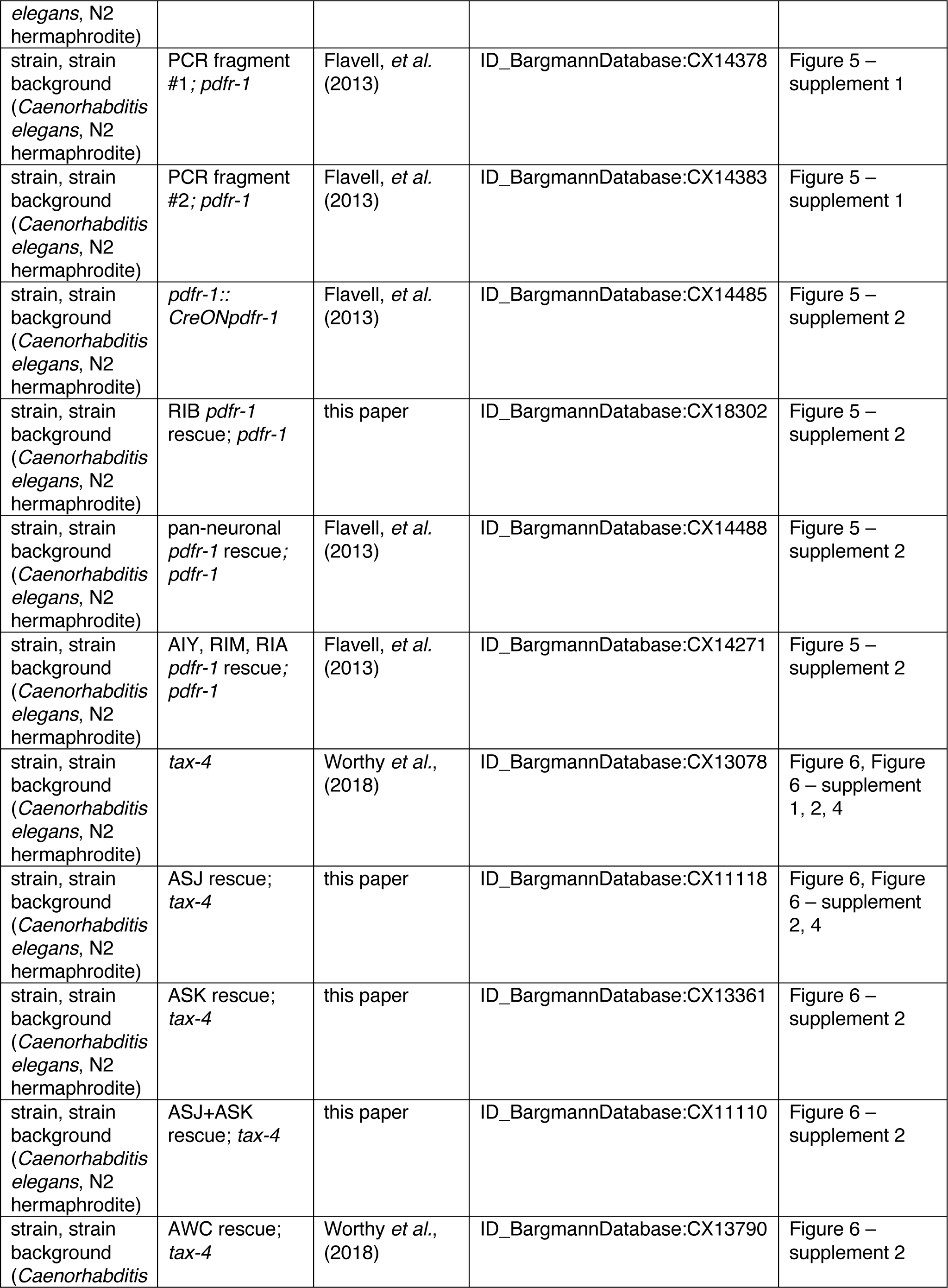
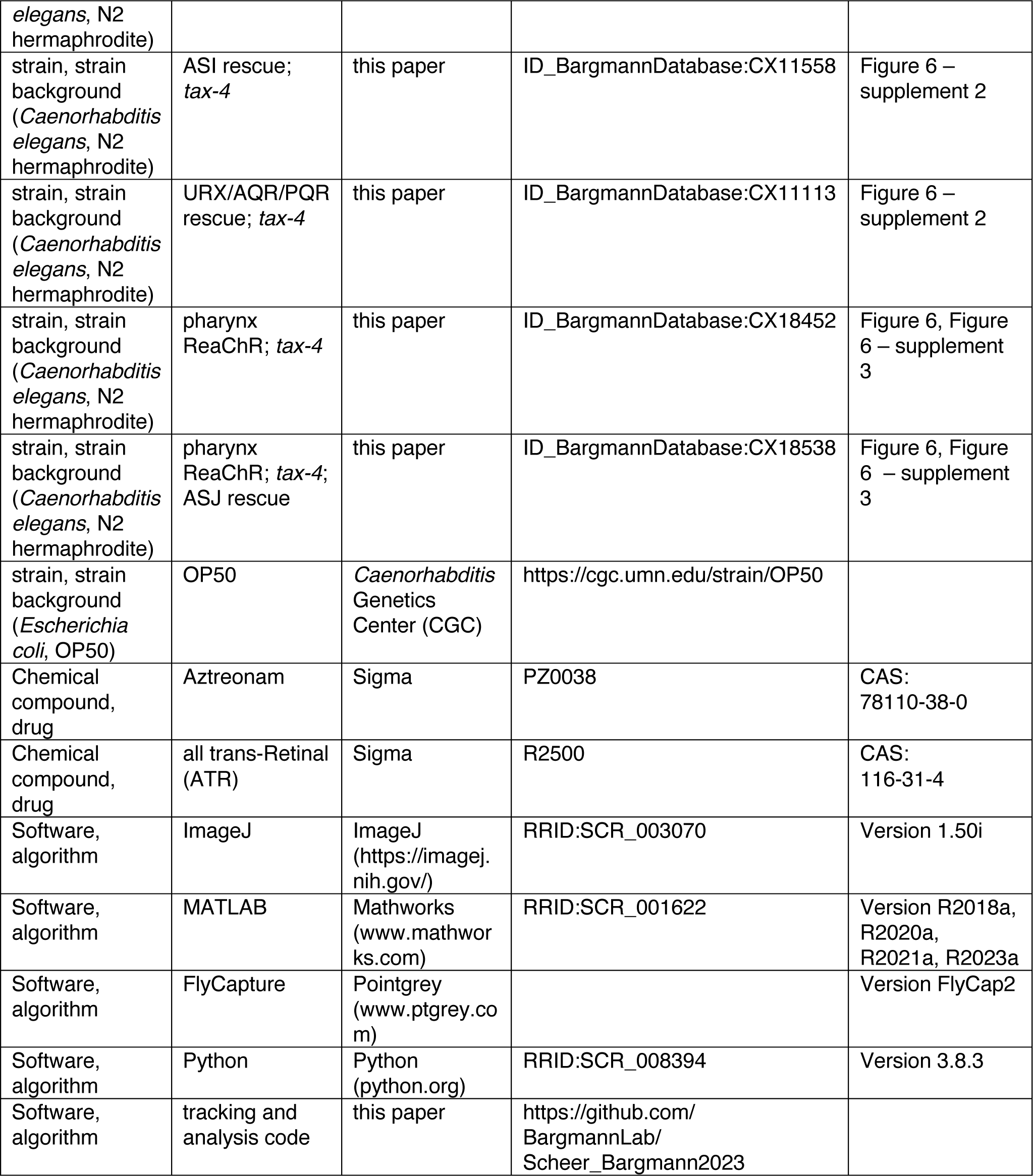

### Nematode and bacterial culture

Bacterial food used in all experiments was *E. coli* strain OP50. Nematodes were grown at 20°C on nematode growth media plates (NGM; 51.3 mM NaCl, 1.7% agar, 0.25% peptone, 1 mM CaCl2, 12.9 μM cholesterol, 1 mM MgSO_4_, 25 mM KPO4, pH 6) seeded with 200 μL of a saturated *E. coli* liquid culture that had been grown at room temperature for 48 hours or overnight at 37°C (without shaking) from a single colony of OP50 in 100 mL of sterile LB (Brenner, 1974). All experiments were performed on young adult hermaphrodites, picked as L4 larvae the evening before an experiment. Wild type controls were the PD1074 sequenced strain derived from the N2 Bristol strain (Yoshimura et al., 2019), except for Figure 4 *tph-1* and *cat-2* controls, which were the CX0001 isolate of the N2 Bristol strain. Mutant strains were backcrossed into wild type to reduce background mutations. Transgenic strains were always compared to matched controls tested in parallel on the same days. Full genotypes and detailed descriptions of all strains and transgenes appear in Table 3: Strain details.

**Table 3:**
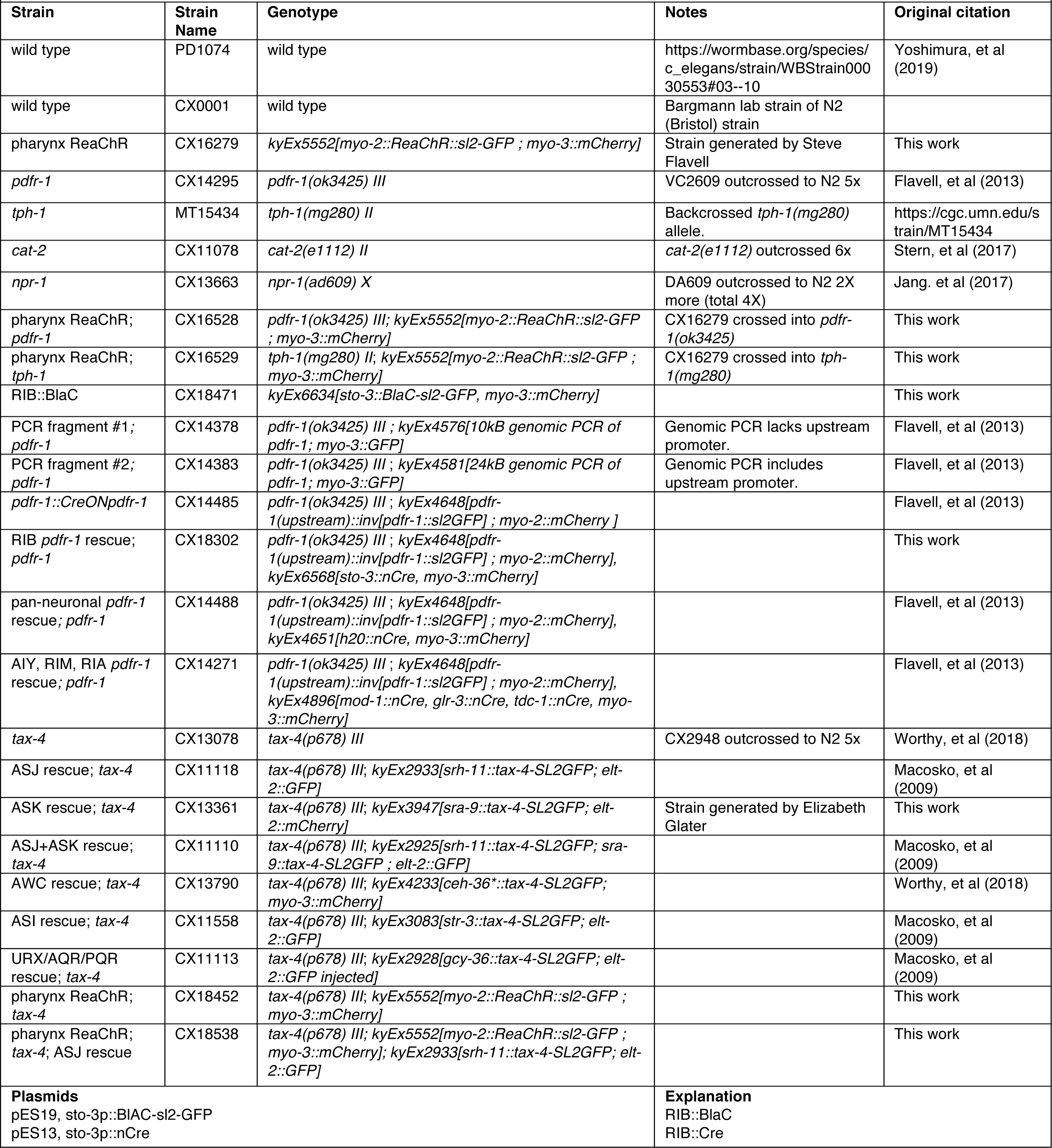
Strains and Plasmids.

### Molecular biology and transgenics

Strains tested for *pdfr-1* rescue using PCR-amplified genomic fragments (Figure 5-S1) were from (Flavell et al., 2013). For cell-selective *pdfr-1* rescue, an inverted cDNA under the *pdfr-1* distal promoter was the floxed rescue construct (Figure 5-S2), and Cre expression was driven by a pan-neuronal promoter (*tag-168*), in RIM (*tdc-1*), in AIY and others *(mod-1*), in RIA *(glr-3),* and in RIB *(sto-3).* A 972 bp region upstream of the *sto-3* gene that drives expression solely in the RIB neurons was cloned using these primers: *sto-3:* gatgcccaatcagttttttttcaccaa, aagccaaaccaagtgagaagaagtattca

Strains and extrachromosomal arrays for *tax-4* rescue were reported in Macosko et al., 2009 and Worthy et al., 2018. The sequences of the promoter ends are given below, along with the concentration at which the plasmid was injected for generating transgenic lines.

ASJ *tax-4* rescue

*srh-11*: gggcaaggacaatgttgccgcag, tgggaataaaataacgacgtatgaata, 50 ng/μl

ASK *tax-4* rescue

*sra-9*: gcatgctatattccaccaaaagaaa, tagcttgtgcatcaatcatagaaca 50 ng/μl

AWC *tax-4* rescue

*ceh-36*: ctcacatccatctttctggcgact, ttgtgcatgcgggggcaggcga, 30 ng/μl

ASI *tax-4* rescue

*str-3*: gtgaacttgaaaagcgcaagtgatat, ttccttttgaaattgaggcagttgtc, 100 ng/μl

URX/AQR/PQR *tax-4* rescue

*gcy-36*: tggatgttggtagatggggtttgga, aaattcaaacaagggctacccaaca 2 ng/μl

Transgenic animals were generated by microinjection of DNA containing the genetic construct of interest, a fluorescent co-injection marker (*myo-2p*::mCherry, *myo-3p*::mCherry, *elt-2p*::nGFP, *elt-2p*::mCherry), and empty pSM vector to reach a final DNA concentration of 100 ng/μL. Transgenes were maintained as extrachromosomal arrays.

### Small lawn foraging assay

For all assays, *E. coli* OP50 was grown overnight in a shaking LB liquid culture from a single colony at 37°C. On the morning of the assay, 400 μL of saturated liquid culture was diluted into 5 mL of LB and allowed to grow to OD1 at 37 °C (∼1.5 hours), as measured by spectrophotometer. The liquid culture was then spun down and resuspended in M9 buffer (3 g KH_2_PO_4_, 6 g Na_2_HPO_4_, 5 g NaCl, 1 ml 1 M MgSO_4_, H_2_O to 1 liter) then concentrated to a density of OD2 (and OD1 or OD4 in Figure 3A-B). To generate the test lawns, 2 μL of this concentrated bacterial resuspension was seeded onto NGM agar in the center of each well of a custom-made laser-cut 6-well plate, where each well is 10mm in diameter (Stern et al., 2017).

50 μL of bacterial resuspension was seeded onto a separate NGM agar plate to be used as a food density acclimation plate. Lawns were grown at 20-22°C for 2 hours before the assay. Adult hermaphrodites picked as L4s 16-20 hours before the assay were then transferred to acclimation plates. After 45-90 minutes, animals were transferred to an unseeded NGM plate, cleaned of *E. coli*, and transferred singly into each well of the assay plates, where they were placed on bacteria-free agar and allowed to find the small food lawn on their own. Animals of the same genotype were grouped on the same 6-well plates and each plate was recorded by a single camera. We used 12 cameras, enabling simultaneous recording of up to 72 individual animals at a time. Temperature and relative humidity within the behavioral recording apparatus were continuously monitored during recordings to ensure that environmental conditions were consistent across filming locations. As a further precaution, the filming locations of each genotype and wild type controls within the recording apparatus were randomized across batches of experiments and days to average out behavioral influences deriving from non-uniform local environmental conditions. Assays were recorded for 1 hour at 3 frames per second using 12 8.8 MP USB3 cameras (Pointgrey, Flea3) and 35 mm high-resolution objectives (Edmund Optics). LED backlights (Metaphase Technologies) provided uniform illumination of the assay plates. Commercial software (Flycapture, Pointgrey) was used to record the movies.

### Uniform lawn assay

Assays testing worm behavior on uniform bacterial lawns were performed as in the small lawn foraging assay with the exception that instead of 2 μL of bacteria seeded in the center of the well, 15 μL of OD2 bacterial suspension was spread evenly throughout the well. Bacteria were grown for 2 hours before acclimation and starting the assay.

### Optogenetic feeding inhibition assay

Pharyngeal pumping was inhibited by expressing the red-shifted channelrhodopsin ReaChR (Lin et al., 2013) under the *myo-2* pharyngeal muscle promoter (a strain generously provided by Steve Flavell). Animals were stimulated while navigating small food lawns described above. Experimental animals were grown on bacterial lawns containing 50 μM all-trans retinal (+ATR) overnight before assays. Control animals were placed on lawns made in parallel that did not contain retinal (-ATR). A 590 nm Precision LED with Uniform Illumination (Mightex) controlled with custom MATLAB software was used to deliver optogenetic stimuli. Animals were acclimated to small lawns (no ATR) for 20 minutes before being exposed to alternating 10-minute intervals of light OFF and light ON using 590 nm light at 60 μW/mm^2^ strobed at 10 Hz with a 50% duty cycle (Fig. 3, Fig 4-S1, Fig. 6, Fig 6-S3). Recording hardware and software was identical to that of off-food foraging assays without optogenetic stimulation except that 475 nm short-pass filters were used on recording optics (Edmund Optics) to prevent overexposure of the video recording during light pulse delivery.

For behavioral quantification in Fig. 3E-F, Fig. 4-S1A,C-D, and Fig 6-S3A, only data before, during and after the first light pulse is shown. For aggregate comparisons of within-animal behavior in light OFF vs. light ON conditions, the two light OFF and light ON pulses were merged, i.e. merging intervals 0-10 min + 20-30 min, and 10-20 min + 30-40 min for statistical analyses. For statistical tests comparing the fraction of animals roaming across genotypes or conditions, steady state light intervals were used for averaging and comparing as indicated in the figure legends: 0-10, 12-20, 22-30 minutes.

### RIB::BlaC assay

Activation of the RIB neurons was accomplished using blue light-activated adenylyl cyclase BlaC (Ryu et al., 2010). BlaC was cloned into the pSM vector under the *sto-3* promoter to drive expression in the RIB neurons. Stimulation was carried out as described for ReaChR (above), except no all-trans retinal was applied and the animals were exposed to blue light (455 nm) at 3 μW/mm_2_ strobed at 10 Hz with a 50% duty cycle (Fig. 5, Fig 5-S3). 525 nm long pass filters were used on recording optics (Edmund Optics) to prevent overexposure of the video recording during light pulse delivery. Behavioral quantification for Fig. 5B-C was conducted as above (optogenetic feeding inhibition assay).

### Inedible food assay

Experiments with inedible bacterial food generated by addition of aztreonam were performed following the protocol of (Gruninger et al., 2008). *E. coli* OP50 was grown in LB from a single colony to saturation overnight. On the morning of the assay, 400 μL of saturated liquid culture was diluted into 5 mL of LB with aztreonam (10 μg/ml) and allowed to grow to OD1 at 37 °C. To generate test lawns, 2 μL of this bacterial suspension was plated on NGM agar test plates containing aztreonam (10 μg/ml). Small lawn assays were performed as described above (see “Small lawn foraging assay”), except that bacteria on NGM+aztreonam plates were allowed to grow for 4 hours before assay testing. To test for any acute behavioral responses to aztreonam, we also performed a control “post-add” experiment, in which 2 μL of either 4 μg/mL aztreonam dissolved in DMSO or DMSO alone was added to normal small bacterial lawns after 4 hours of growth on NGM agar (Fig 3-S1).

### Behavioral tracking and lawn feature detection

Because each video recorded the behavior of up to 6 individual animals, videos were manually cropped so each surrounded just a single animal using FFmpeg software (Tomar, 2006). To extract animal positions and postures, captured movies were analyzed by custom made scripts in MATLAB (Mathworks, version 2021a) using the Image Processing Toolbox and the Computer Vision Toolbox. In each frame of the movie, the worm is segmented by background subtraction and its XY position is tracked over time using a Kalman filter. From the background-subtracted worm image, a smooth spline of 49 points was computationally applied and features relating to the movement of points along the body were derived following (Javer et al., 2018). Disambiguation of the head versus tail was determined by assigning the head as the end of the spline that had greater cumulative displacement over the video assay, facilitating determination of times when the animal moved forward and backward.

#### Behavioral features extracted

Features were defined as described in (Javer et al., 2018). Briefly, body parts were defined based on a skeletonized spline containing 49 points equally distributed along the length of the worm body. The head comprises spline points 1-8. The midbody comprises spline points 17-33. The positions of head and midbody were calculated by deriving the centroid of each of these point sets by averaging their x and y positions before subsequent analyses (see Fig 2-S2).

- *Midbody speed:* the derivative of displacement of the midbody across frames. Positive and negative numbers indicate forward and backward motion, respectively. Units: mm/sec.
- *Midbody angular speed:* The angle change across midbody positions over time: Two vectors are measured: *v_0-1_*, representing the change in midbody position from time frame 0 to 1, and *v_1-2_*, representing the change in midbody position from time frame 1 to 2. the position change over three frames at each time point was quantified. Angular speed is defined as the arc-cosine of the dot product of *v_0-1_* and *v_1-2_* divided by the scalar product of the norms of these vectors. Units: degrees per second.
- *Head speed:* the derivative of displacement of the head across frames. Units: degrees per second.
- *Head angular speed:* Same as midbody angular speed but calculated for the head position. Units: degrees per second.
- *Head radial velocity relative to the midbody*: The derivative of displacement of the head relative to the midbody across frames. This is calculated by subtracting the midbody position from the head position and shifting to polar coordinates (*Φ*,*r*) where *Φ* is the angular dimension and *r* is the radial dimension. Head radial velocity relative to midbody is the derivative of *r* with respect to time. Units: mm/sec.
- *Head angular velocity relative to the midbody:* the derivative of the angular displacement of the head relative to the midbody across frames. The same procedure as above is used to generate polar coordinates. Head angular velocity relative to the midbody is the derivative of *Φ* with respect to time. Units: degrees per second.
- *Quirkiness:* 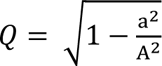, where a is the minor and A is the major axis of a bounding box surrounding the animal, respectively. Values closer to 1 means the animal’s shape is more elongated and thinner; closer to 0 indicates a more rounded shape.

All of these features were binned into contiguous 10-second intervals for subsequent analyses. There are also several features only defined for 10-second bins:

- *Fraction of time moving forward* per bin.
- *Fraction of time moving reverse* per bin.
- *Fraction of time paused* per bin.
- *Midbody forward speed*, the mean of Midbody speed where values are >= 0.
- *Midbody reverse speed*, the mean of Midbody speed where values are <= 0.

#### Lawn boundary-related metrics and Lawn boundary interaction behaviors

The outline of the bacterial lawn was determined by edge detectors applied to the background averaged across movie frames. Across every frame, the closest boundary point to the animal’s head was determined and used to calculate “lawn boundary distance.”

Lawn boundary distance and movement direction were used to classify a set of lawn boundary interaction behaviors. Head pokes were classified based on an excursion of the head that peaks outside the lawn before returning to the lawn interior. In the period following maximal displacement outside the lawn and before resuming locomotion inside the lawn (“recovery interval”), 3 types of head pokes were categorized: head poke forward, in which at least half of the recovery intervals is spent moving forward, head poke reversal, in which the animal executes a reversal during the recovery interval, or head poke pauses, in which an animal spends at least half of the recovery interval with speed less than 0.02 mm/sec. Lawn leaving events were marked as the first frame when the animal’s head emerged from the lawn before its entire body exited the lawn.

### Quality control for including animals in subsequent analyses

Behavior and lawn features were detected and tracked over the 1-hour assay but only the latter 40 minutes of data were retained for analysis to minimize the effects from manipulating animals prior to recordings. Data from single animals were only retained in subsequent analyses if the following conditions were met: 1) the worm was visible in the video for at least half of the time the worm was recorded (cumulatively 30 minutes), 2) the worm was inside the bacterial lawn for at least one minute within the first 20 minutes of the video (before usable data was collected), 3) the plate was not bumped during the recording. All criteria were established before data collection.

### Quantitative Locomotion Analysis and HMM analyses

All quantitative analyses of locomotion and Hidden Markov Model building after behavioral tracking were performed in Python. For all analyses of animal locomotion and model-building, behavioral data from animals outside the food lawn was censored.

To classify roaming and dwelling states, speed and angular speed of animal centroid position was averaged into contiguous 10 second intervals. Roaming and dwelling states were classified as in (Flavell et al., 2013). Briefly, two classes of intervals corresponding to high angular speed / low speed and low angular speed / high speed were identified and separated by a line drawn at y (mm/sec) = x (deg/sec) /450. Behavior can then be instantaneously classified into roaming intervals when values are above the line, or dwelling intervals, when values are below the line. A 2-state categorical Hidden Markov Model was then trained on these behavioral sequences to generate roaming and dwelling hidden states using the SSM package (Linderman et al., 2020).

An Autoregressive Hidden Markov Model (AR-HMM) was trained to segment animal behavioral states on food using a different set of behavioral features relating to forward body movement and head movements: fraction of time moving forward per 10-sec bin, midbody forward speed, midbody angular speed, head angular velocity relative to midbody and head radial velocity relative to midbody (Fig. 2, Fig 2-S2, see above for feature definitions).

Formally, at each time step *t*, we have discrete hidden states *z_t_* ∊ 1, 2,…*K* that follow Markovian dynamics, 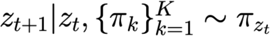, where 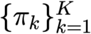 is the transition matrix and *π_k_* ∊ [0, 1]^*K*^ corresponds to the *k*-th row. Given hidden states *z_t_*, the resulting feature dynamics are given by 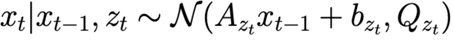, where *A_k_* and *Q_k_* are real 5×5 linear dynamics and covariance matrices, respectively. *b_k_* ∊ ℝ^5^ is the bias. The linear dynamics matrix *A* specifies a continuous flow on the feature space. The bias term *b* can drive the dynamics in a particular direction. In the case where *A* is all zeros, the system has no dynamics and this amounts to a Hidden Markov Model with Gaussian emissions. AR-HMMs were also trained using the SSM package (Linderman et al., 2020). AR-HMM performance was evaluated by calculating the ratio of the log-likelihood of held-out test data set using the AR-HMM same data under the AR-HMM and a multivariate Gaussian model 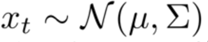.

### Sample Size Determination, Replicates and Group Allocation

The number of animals in each experiment is detailed in the figure legends. Assays were typically repeated across at least two days of experiments to account for day-to-day variability. Control animals were always run on the same days as experimental animals.

Using G*POWER 3.1, we chose sample sizes based on the desired ability to detect an effect size of 1 with 80% power and a 5% alpha, yielding a minimum value of n=18 animals per group for a two-sample unmatched comparison of means and n=11 animals per group for a matched comparison of means (Faul et al., 2009).

### Quantification and Statistical Analysis

All comparisons in the fraction of time roaming were performed after logit-transformation 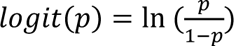, where p is the fraction of time spent roaming per animal.

Either Wilcoxon rank sum test or paired t-tests were used to compare matched data (roaming speed at −3 to −1 minutes vs. last 30 seconds before leaving, number of lawn leaving events during light OFF vs. ON, fraction of time roaming during light OFF vs. ON).

The Kolmogorov-Smirnov test was used for comparing cumulative distributions.

If multiple pairwise tests were done, multiple hypothesis correction was always performed.

Statistical details for each experiment are described in the figure legends. The p values resulting from all statistical tests performed in the paper can be found in Table 4.

**Table 4:**
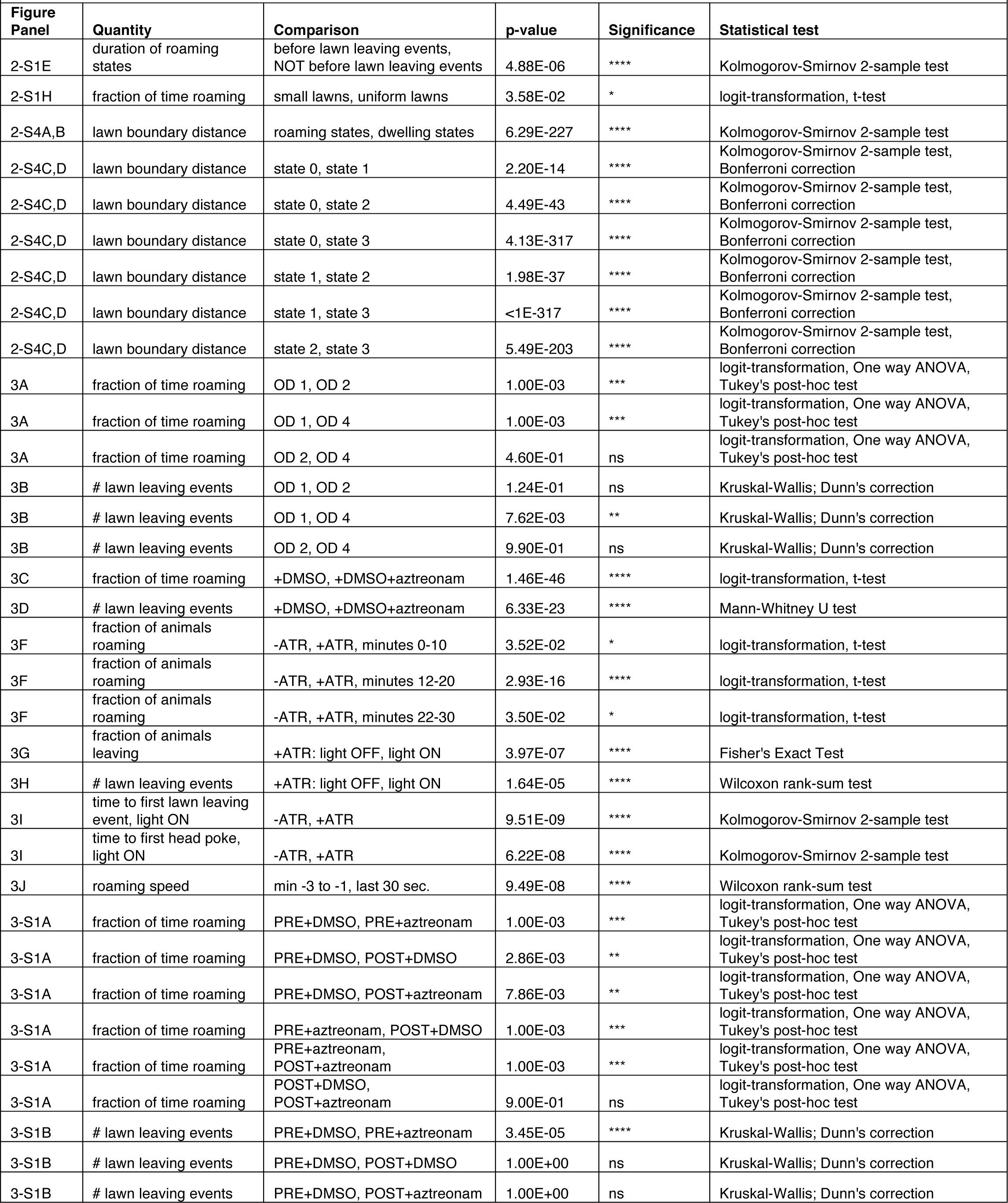
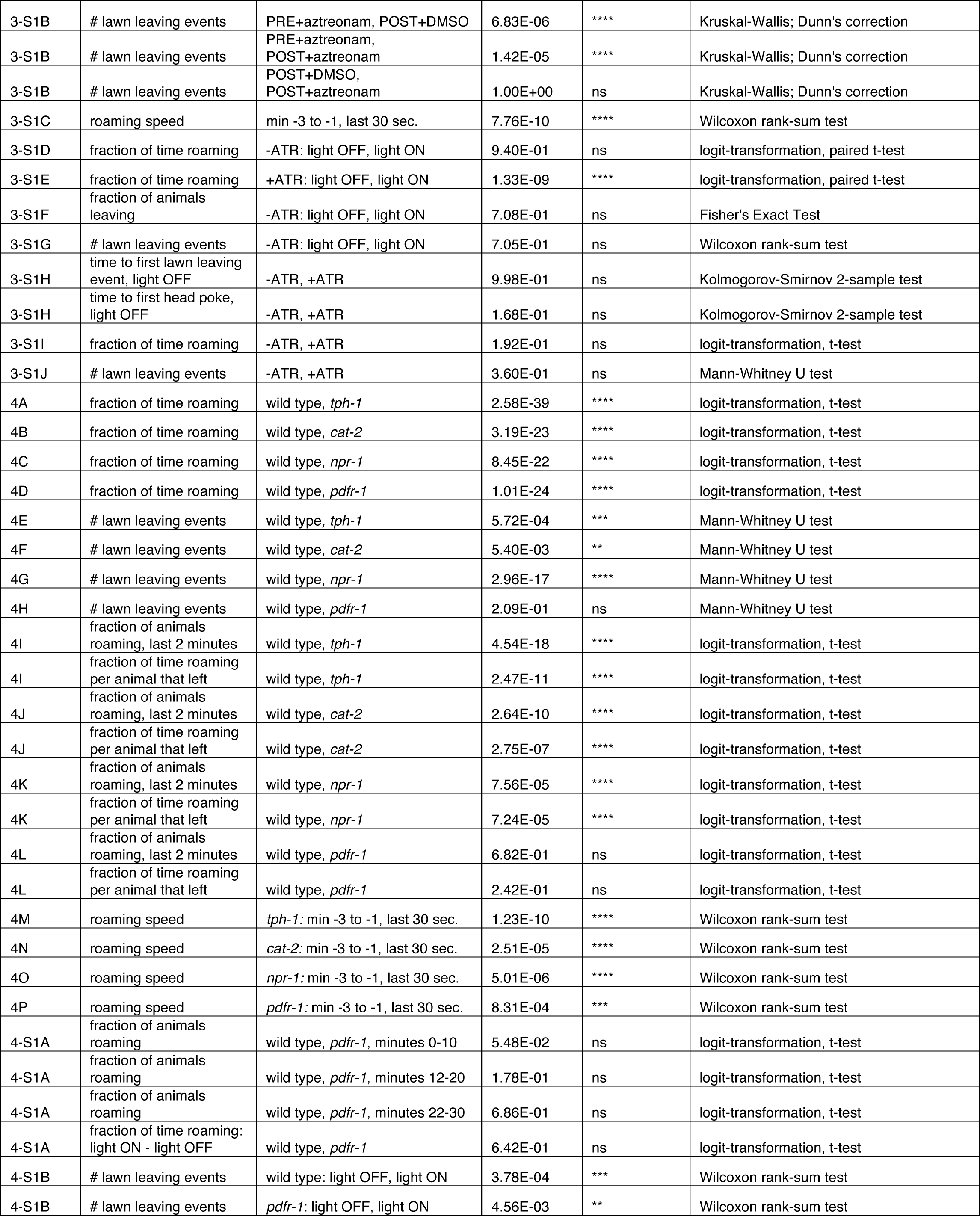
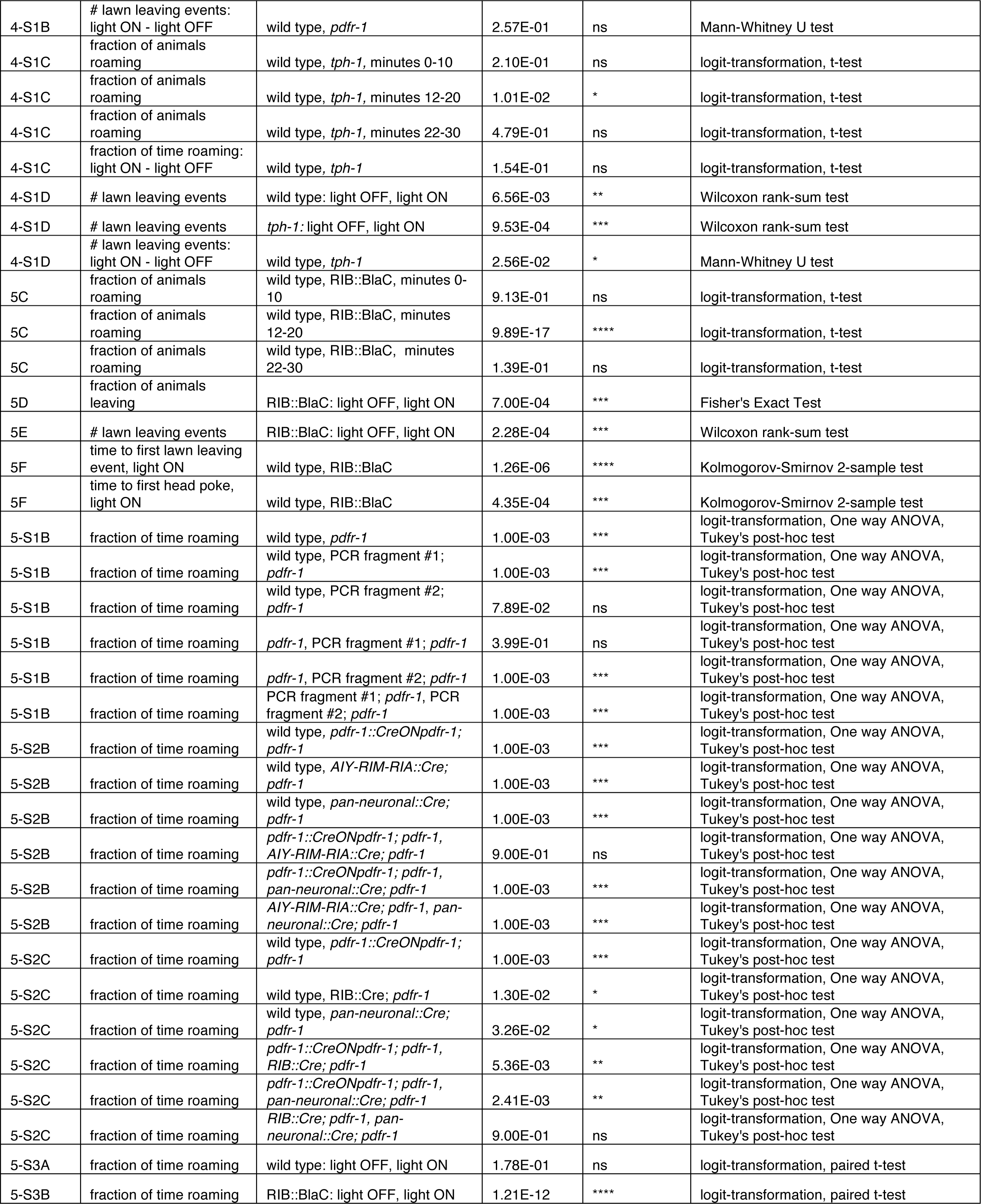
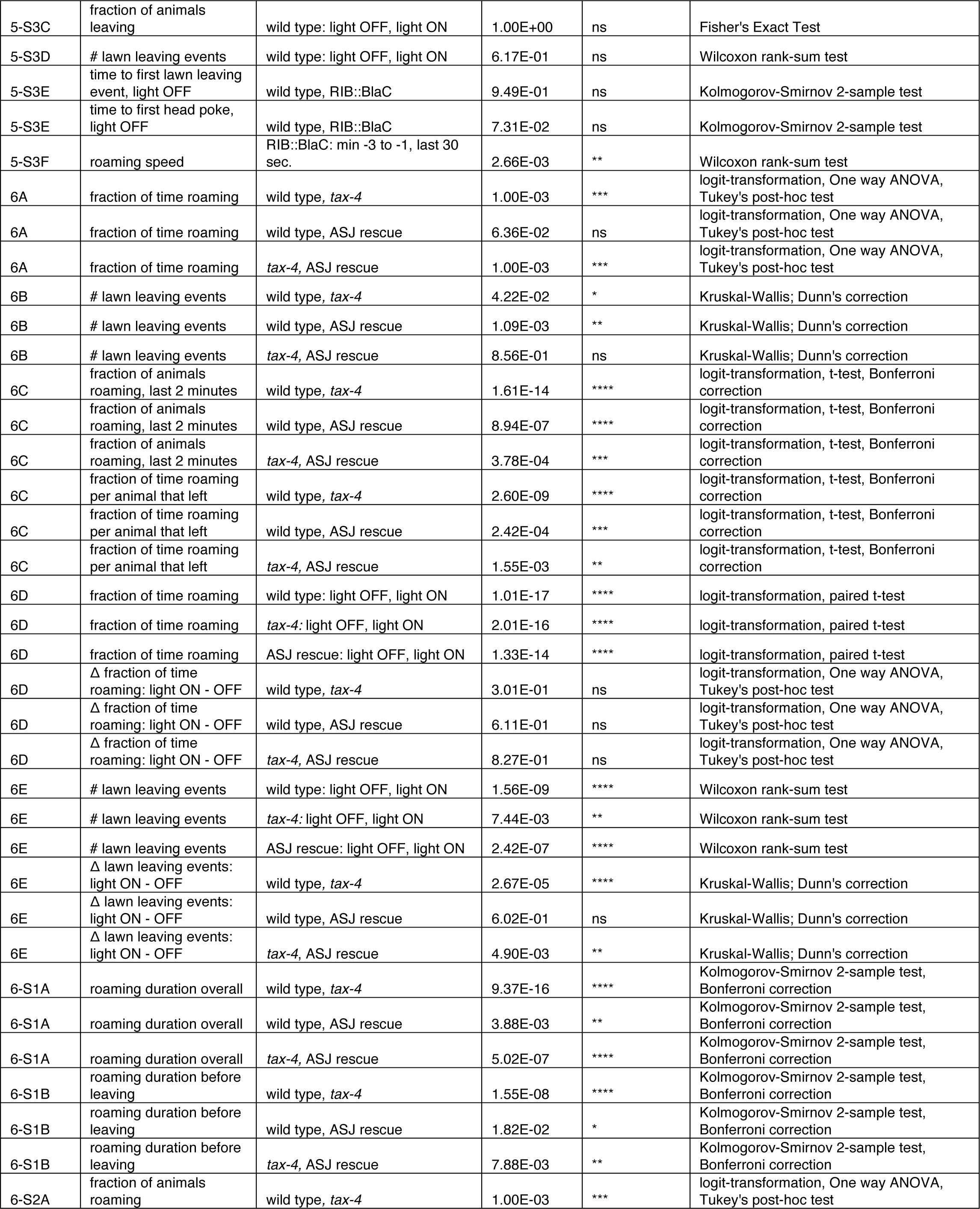
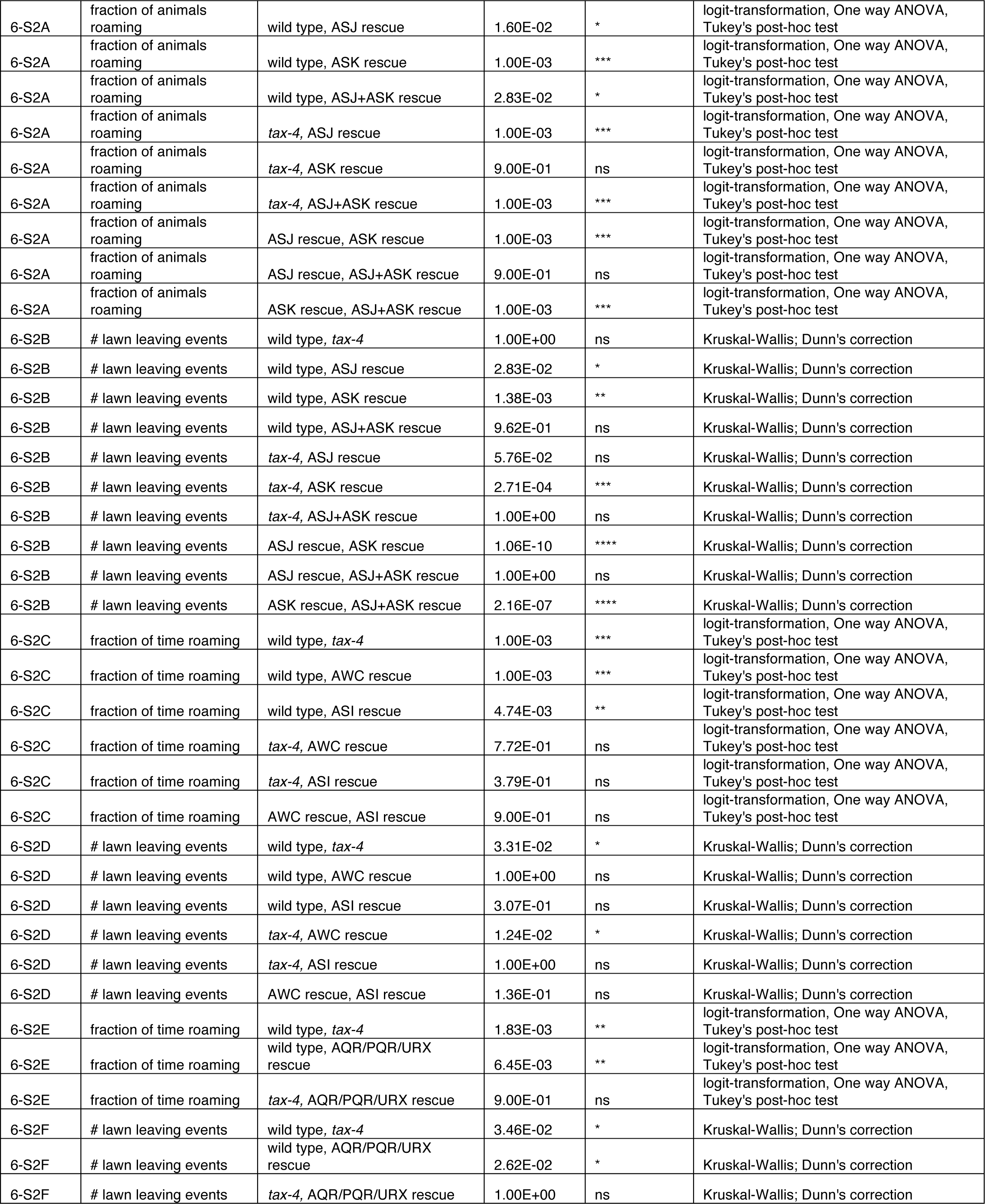
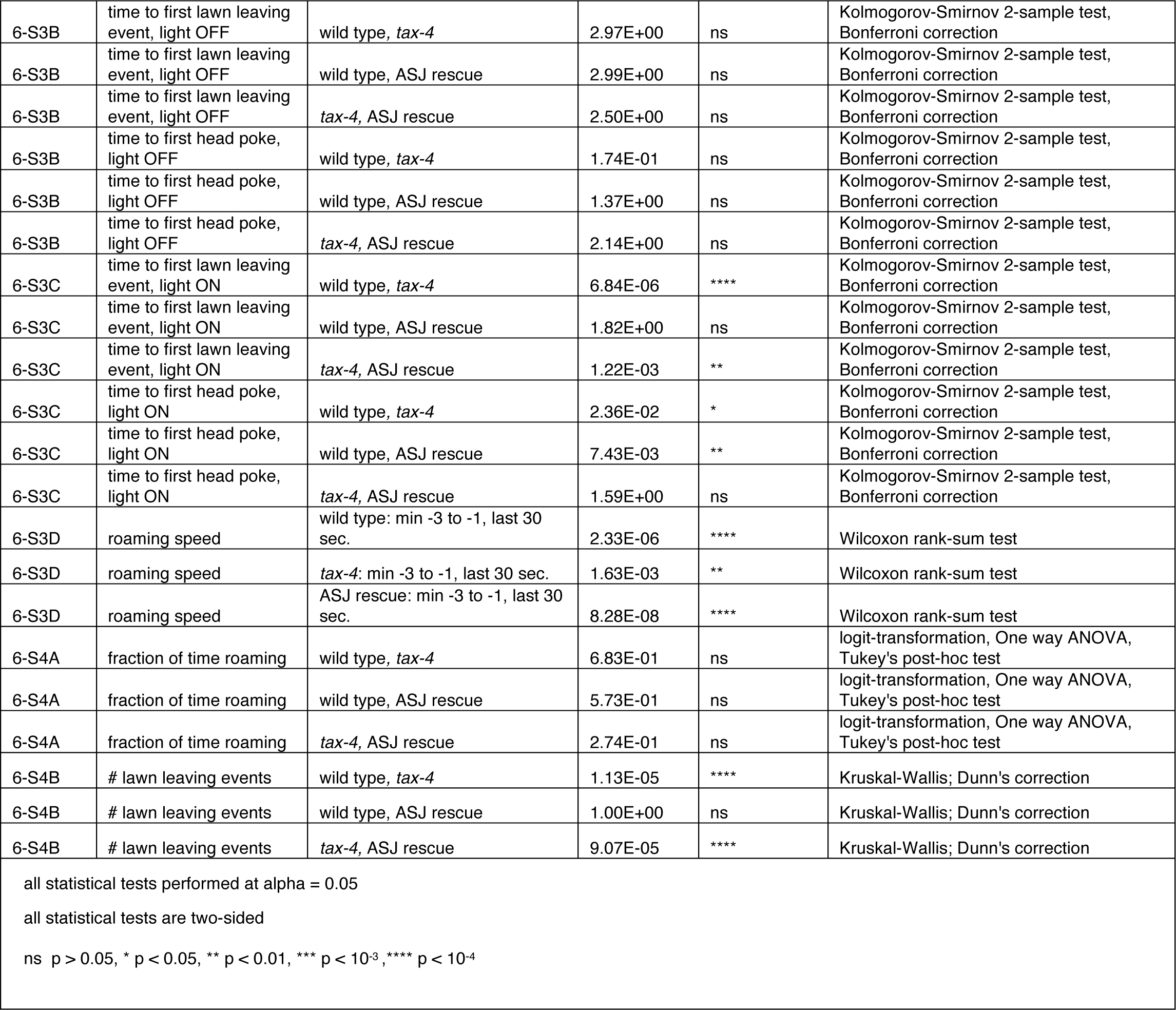
p-values.

### Materials and Data Availability Statement

All primary behavioral data and relevant code for data analysis are available at Dryad (https://doi.org/10.5061/dryad.47d7wm3jf) and Github (https://github.com/BargmannLab/Scheer_Bargmann2023) without restriction. Source data files contain the summarized data for all plots in Figures 1-6.

All strains and plasmids used are found in Table 3: Strain details and are available from the corresponding author without restrictions.

## ACKNOWLEDGMENTS

We thank Cheryl Mai for initial analysis of optogenetics experiments. We thank Philip Kidd, Audrey Harnagel, Yarden Wiesenfeld, and Steven Flavell for thoughtful discussions and comments on the manuscript. This work was supported by the Chan Zuckerberg Initiative and by an NIH F31 Predoctoral NRSA Fellowship to E.S.

## Supplemental Videos

**Figure 1 – supplement video 1:**

Lawn boundary encounters lead to different behavioral responses: lawn leaving, head poke forward, head poke pause, and head poke reversal. Green dot indicates the head position. White line indicates the lawn boundary. Forward and Reverse motion indicated in top left corner.

**Figure 2 – supplement video 1:**

Example of animal behavior with roaming and dwelling annotated. Animal shown corresponds to the behavior before and after the first lawn leaving event in Fig. 2-S2P.

Green dot indicates the head position. White line indicates the lawn boundary. Forward and Reverse motion indicated in top left corner. Speed indicated top middle. Time stamps indicated top right. Video sped up 7x.

**Figure 2 – supplement video 2:**

The same animal and time steps as in Fig. 2-Supplemental video 1, except annotated with AR-HMM states instead of roaming and dwelling.

---

**Figure 1 – supplement 1:**
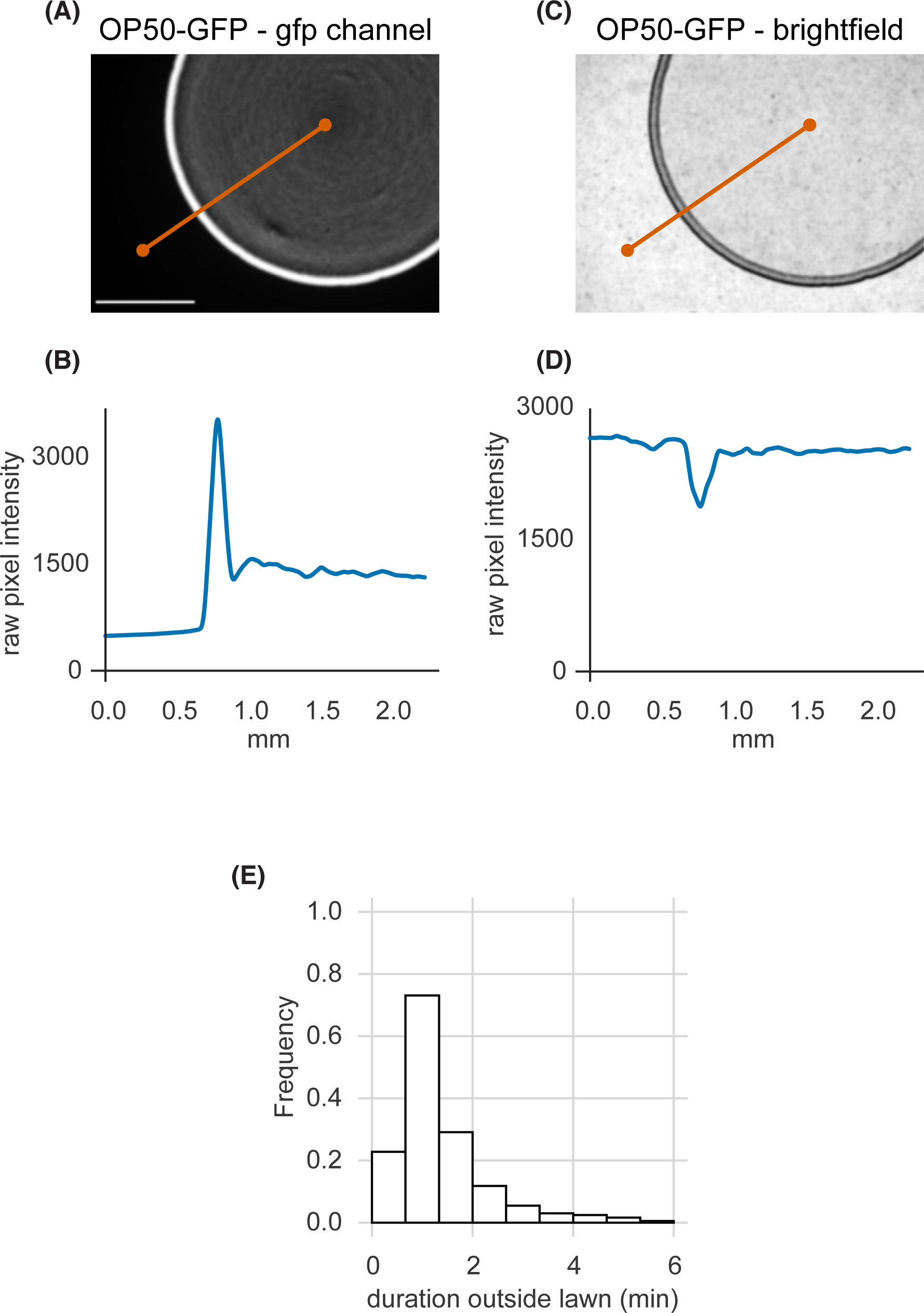
Small bacterial lawns are denser at the lawn boundary, and quantification of duration outside the lawn. **(A)** A small lawn of bacteria expressing green fluorescent protein (GFP) imaged in the GFP channel. White scale bar is 1 mm. **(B)** GFP intensity across the orange transect line plotted in (A) shows accumulation of bacterial cells near the lawn boundary in brighter GFP pixels. **(C)** An image of the same lawn in brightfield. **(D)** Brightfield intensity across the orange transect line plotted in (C) shows local darkening at the lawn boundary corresponding to bright GFP pixels in (A). **(E)** Duration of individual excursions outside the lawn. Quantified across wild type dataset (n = 1586 animals). See Supplementary Table 1.

**Figure 2 – supplement 1:**
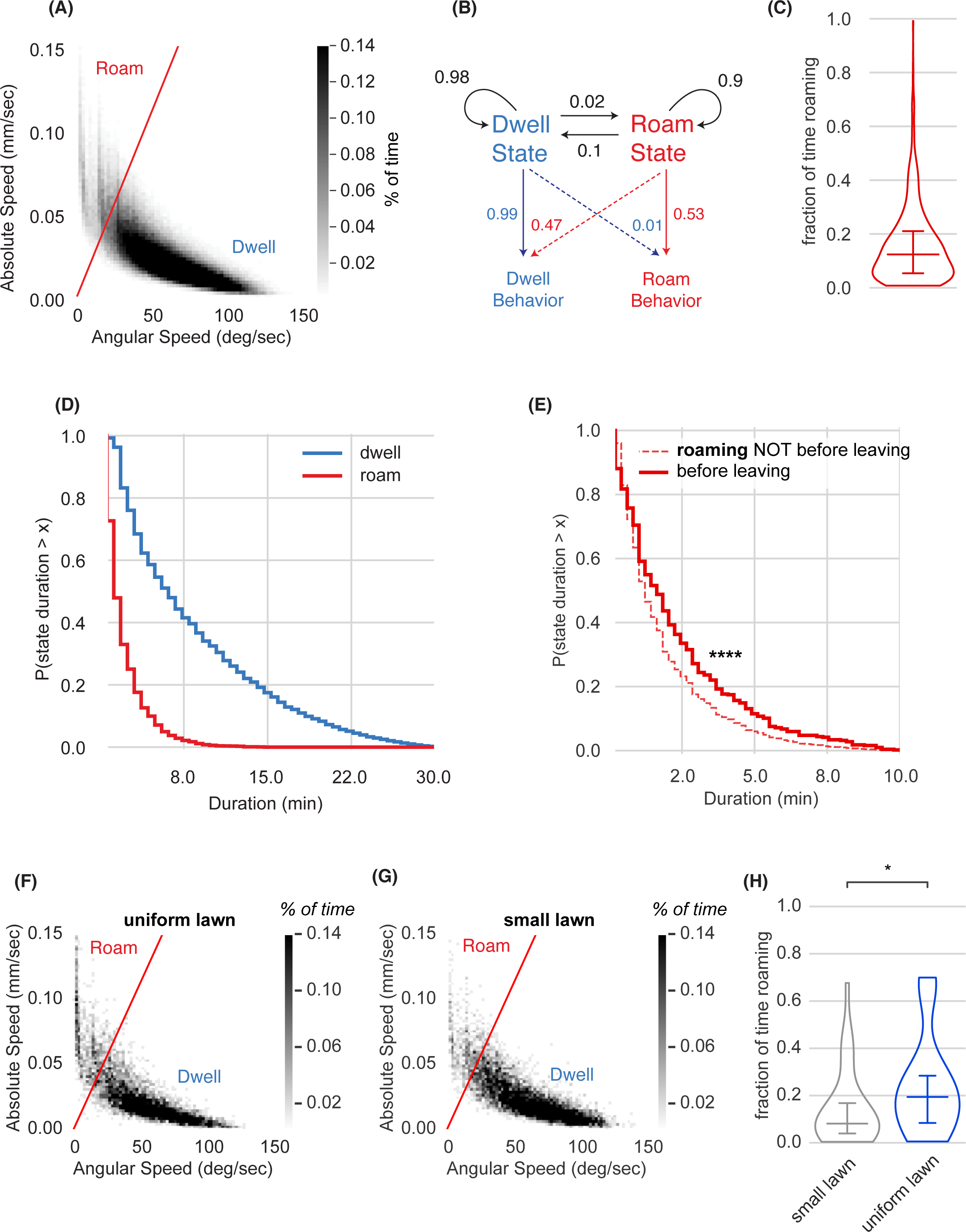
Roaming and dwelling states on small bacterial lawns. **(A)** Scatter plot of average absolute speed and angular speed in 10 second intervals for wild type animals on small lawns (n = 1586 animals, 380,640 bins). The boundary shown, used for all data in the paper, defines roaming (high speed/low angular speed) and dwelling (low speed/high angular speed). **(B)** Two-state Hidden Markov Model trained on wild-type animals. Roaming Behavior and Dwelling Behavior correspond to 10 sec intervals either above or below the separating line in (A), respectively. Roaming and dwelling states are inferred from the Hidden Markov Model. Black arrows indicate transition probabilities between states and colored arrows represent emission probabilities. **(C)** Fraction of time spent roaming across wild type animals. **(D)** Complementary cumulative distribution functions (ccdfs) for roaming and dwelling state durations in minutes. **(E)** Empirical complementary cumulative distribution functions (ccdfs) comparing state durations of roaming states that did or did not precede a lawn leaving event. Statistics by Kolmogorov-Smirnov 2-sample test. **** p < 10^-4^. **(F)** Scatter plot of average absolute speed and angular speed in 10 s intervals for wild type animals on uniform lawns (n = 31 animals, 7,440 bins). **(G)** Same as (E) for animals on small lawns recorded in parallel with animals on uniform lawns (n = 31 animals, 7,440 bins). **(H)** Animals roam more on uniform lawns than on small lawns. * p < 0.05 by Student’s t-test on logit-transformed data. Violin plots show median and interquartile range. See Supplementary Table 2.

**Figure 2 – supplement 2:**
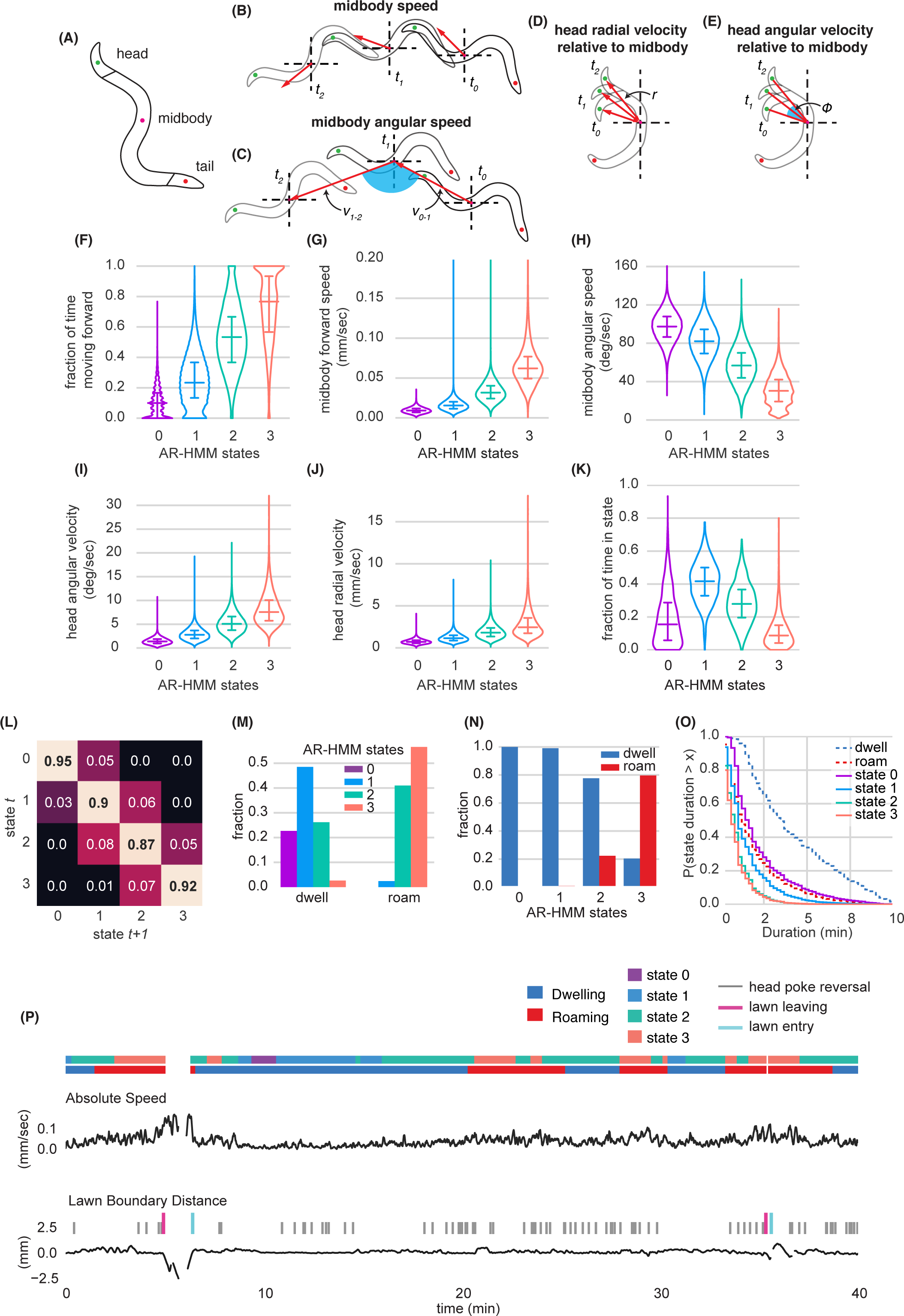
Modeling behavioral states across locomotory feature dimensions using an Autoregressive Hidden Markov Model. **(A)** Schematic of animal body shape with key body points indicated. **(B-E)** Schematics illustrating derived behavioral features. See Methods for detailed description. **(B)** Midbody speed **(C)** Midbody angular speed **(D)** Head radial velocity relative to midbody **(E)** Head angular velocity relative to midbody. **(F-K)** Quantification of behavioral parameters per animal while in each AR-HMM state. **(F)** Fraction of time spent moving forward (G) Midbody forward speed **(H)** Midbody angular speed **(I)** Head angular velocity **(J)** Head radial velocity **(K)** Fraction of time spent in each state per animal. **(L)** Transition matrix for 4-state AR-HMM. Rows index state at time *t*, columns index state at *t+1*. **(M)** Fraction of AR-HMM states found in roaming and dwelling states. **(N)** Fraction of roaming and dwelling states found in each AR-HMM state. **(O)** Empirical complementary cumulative distribution functions (ccdfs) comparing state durations from the 4-state AR-HMM to those of roaming and dwelling. **(P)** Example behavioral trace with two lawn leaving events. Absolute speed and lawn boundary distance are shown for a continuous 40-minute assay. Foraging decisions are annotated with colored tick marks above lawn boundary distance trace. Roaming, Dwelling, and AR-HMM states are annotated above absolute speed with colored bars. Distributions are computed across wild type animals on small lawns (n = 1586). Violin plots show median and interquartile range. See Supplementary Table 2.

**Figure 2 – supplement 3:**
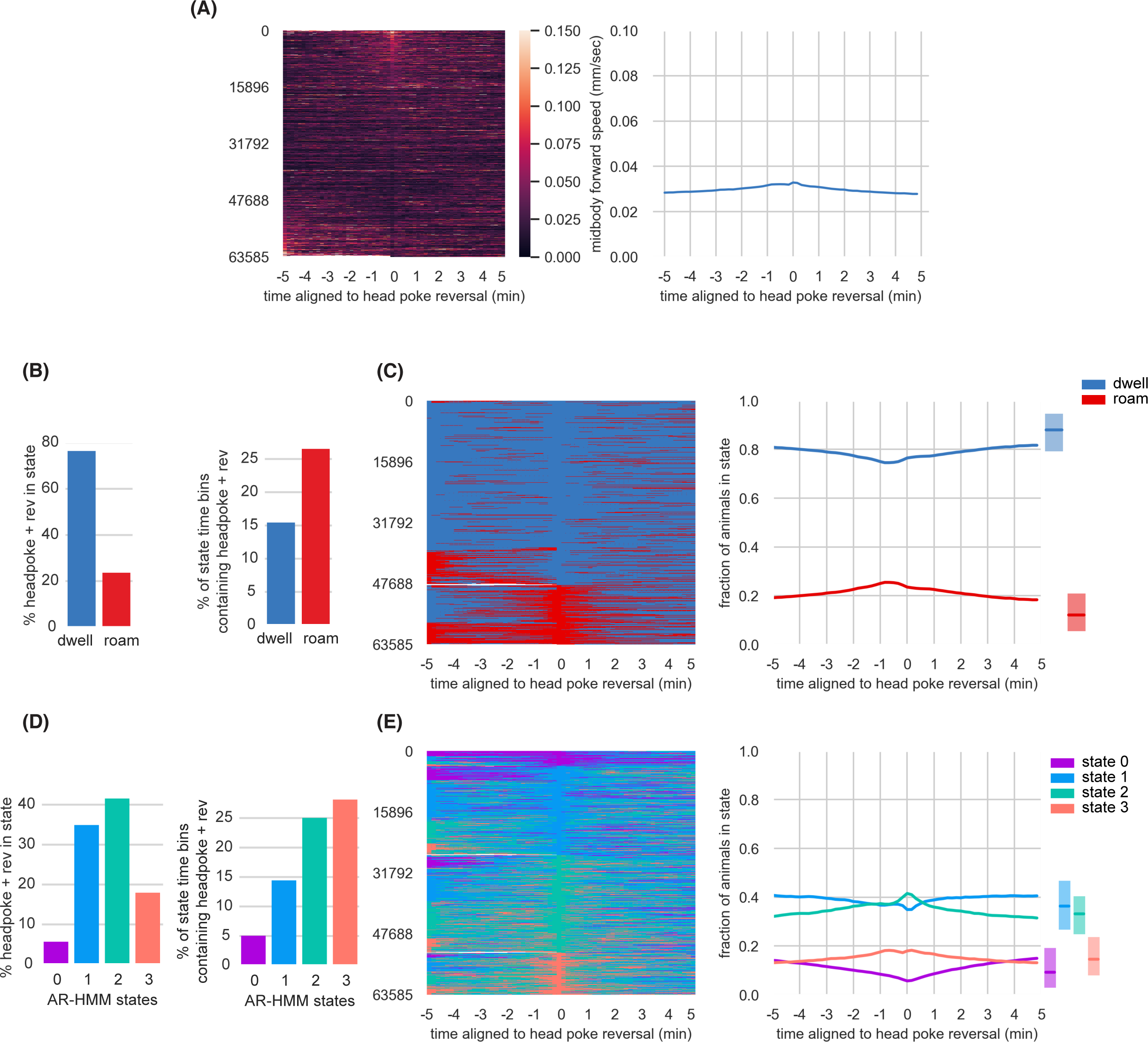
Head Poke Reversals are associated with small changes in arousal states. **(A)** Midbody forward speed aligned to head poke reversals. Left, heatmap of individual speed traces. Right, mean midbody forward speed computed across the heatmap traces. **(B)** Overlap of head poke reversals with roaming and dwelling. Left, percentage of head poke reversals found in each state. Right, percentage of 10 second time bins roaming or dwelling that contain a head poke reversal. **(C)** Roaming and dwelling aligned to head poke reversals. Left, heatmap of roaming/dwelling state classifications aligned to head poke reversals. Center, fraction of animals in roaming or dwelling state prior to head poke reversals. Right, total fraction of time spent roaming and dwelling in all assays that included a head poke reversal. **(D)** Overlap of head poke reversals with AR-HMM states. Left, percentage of head poke reversals found in each state. Right, percentage of 10 second time bins per AR-HMM state that contain a head poke reversal. **(E)** AR-HMM states aligned to head poke reversals. Left, heatmap of roaming/dwelling state classifications aligned to head poke reversals. Center, fraction of animals in each AR-HMM state prior to head poke reversals. Right, total fraction of time spent in each AR-HMM state in all assays that included a head poke reversal. Data aligned to head poke reversals (n = 63,585) from wild type animals (n = 1586). Dark lines represent the mean and shaded region represents the standard error. In boxplots, median is highlighted, and interquartile range is indicated by shaded area. See Supplementary Table 2.

**Figure 2 – supplement 4:**
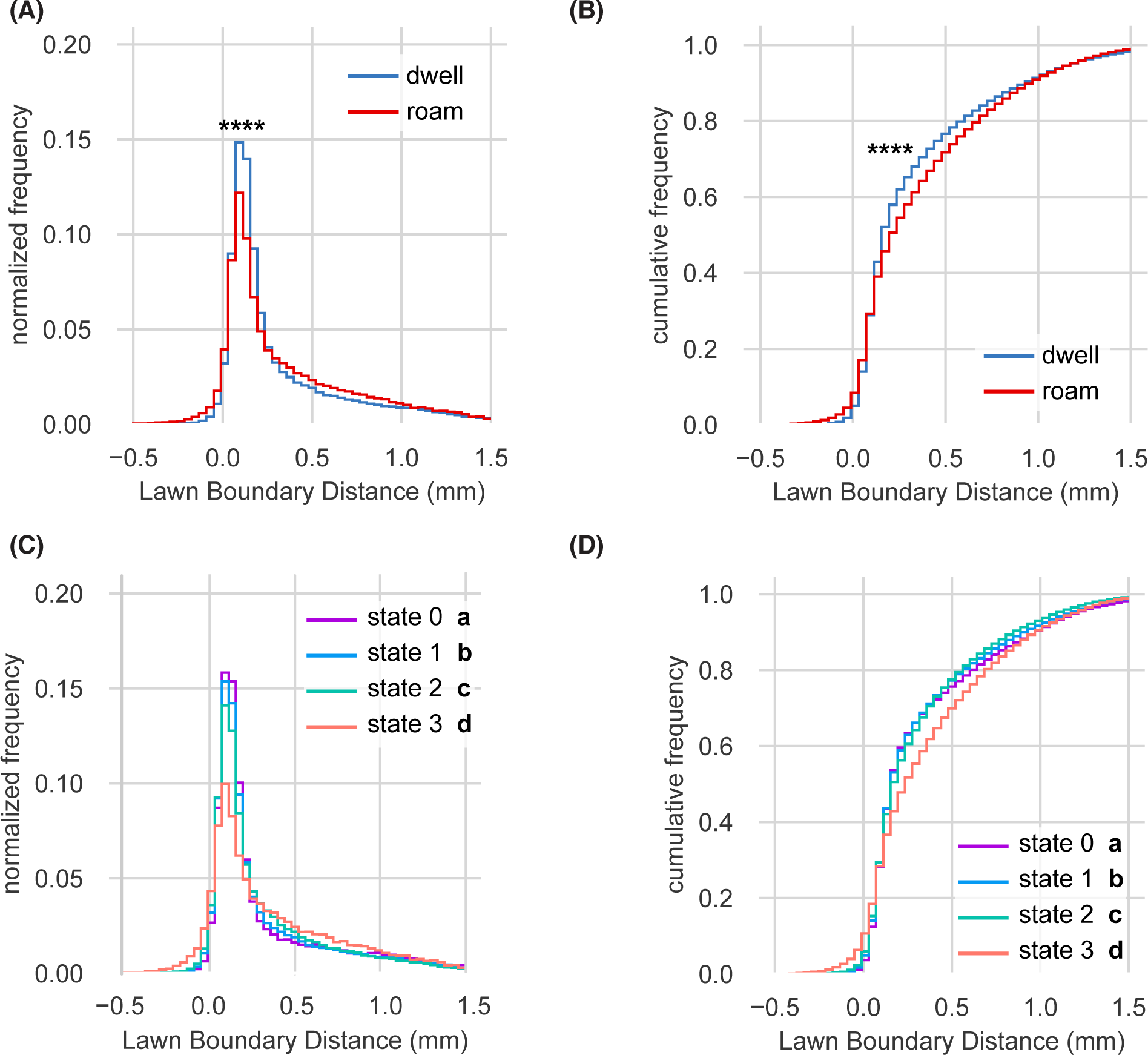
Lawn boundary distance distributions by HMM state. **(A-D)** Empirical distributions of lawn boundary distance by HMM state. Positive values indicate distances inside the lawn, negative values indicate distances outside the lawn (See Fig. 1). **(A)** Lawn boundary distance of animals while roaming and dwelling. Statistics by Kolmogorov-Smirnov 2-sample test. **(B)** Same as (A) but plotted as a cumulative distribution. **(C)** Lawn boundary distance of animals in each state of 4-state AR-HMM. Statistics by Kolmogorov-Smirnov 2-sample tests with Bonferroni correction. **(D)** Same as (C) but plotted as a cumulative distribution. Distributions are computed across wild type animals on small lawns (n = 1586). In **A-B**, **** p < 10^-4^. In **C-D**, each AR-HMM state is significantly different from all others (a-d), although effect sizes are small.

**Figure 3 – supplement 1:**
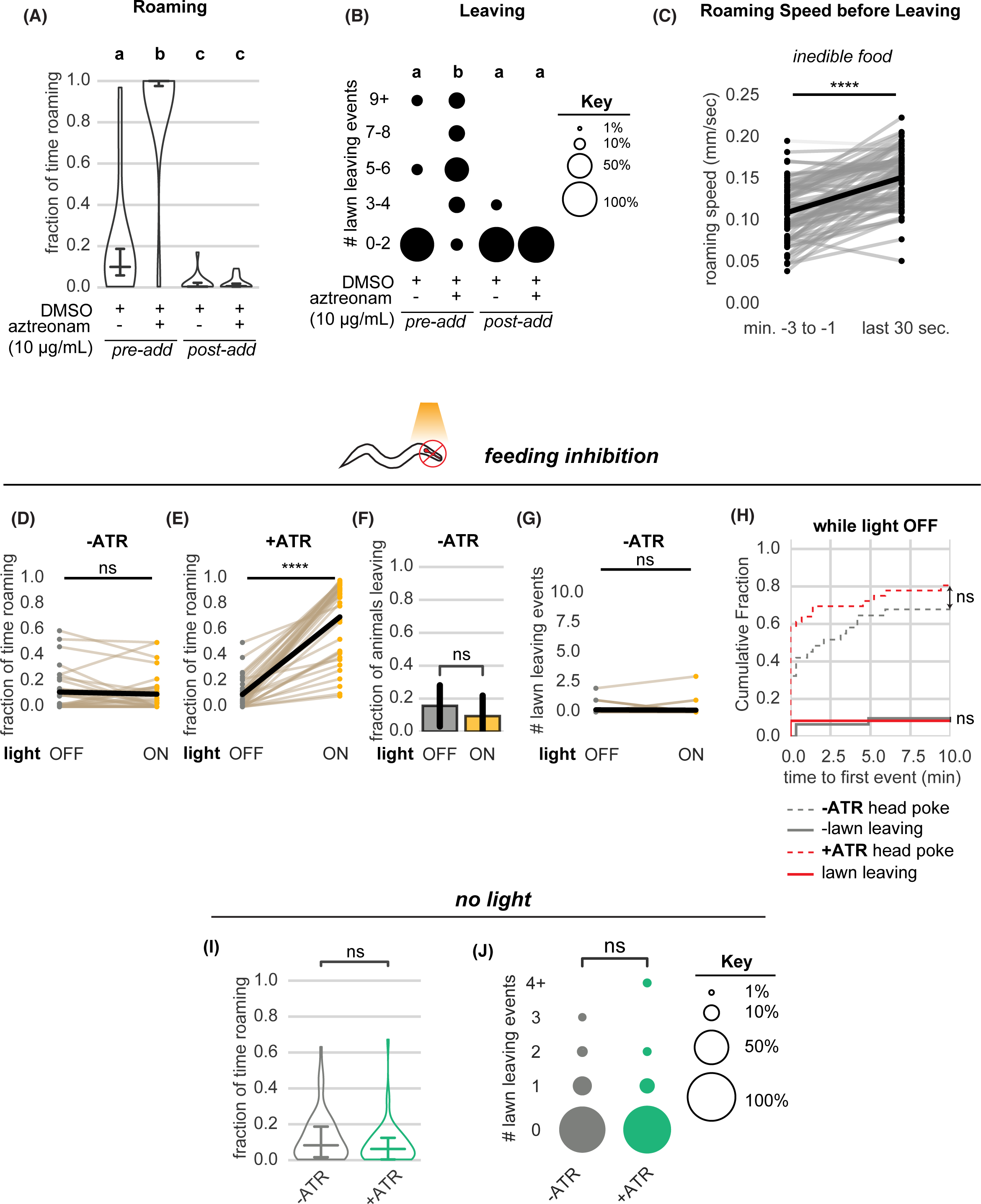
Further quantification of roaming and leaving behaviors under food intake inhibition. **(A-B)** Aztreonam affects roaming and leaving by affecting bacterial growth. In “pre-add” conditions, the drug is added to liquid cultures and agar plates during bacterial growth. In “post-add” conditions, the drug is added after bacterial growth just before testing behavior. **(A)** Fraction of time roaming. Different letters mark significant differences in roaming by logit-transformation followed by one-way ANOVA and Tukey’s post hoc test. (*pre-add*: +DMSO/+aztreonam n = 12, +DMSO (control) n = 15; *post-add*: +DMSO/+aztreonam n = 12, +DMSO (control) n = 16). **(B)** Number of lawn leaving events in the same assays as (A). Different letters mark significant differences in leaving by Kruskal-Wallis followed by Dunn’s multiple comparisons test. **(C)** Animals on inedible food accelerate before leaving the lawn. Statistics by Wilcoxon rank-sum test. **(D-G)** Quantification of controls relating to optogenetic feeding inhibition experiments. **(D)** Light exposure does not induce roaming in animals not pre-treated with all-trans retinal. Statistics by paired t-test on logit-transformed data. **(E)** Light exposure reliably induces roaming in animals pre-treated with all-trans retinal. Statistics by paired t-test on logit-transformed data. **(F)** Light exposure does not increase the fraction of animals that leave lawns when animals are not pre-treated with all-trans retinal. Statistics by Fisher’s exact test. **(G)** Light exposure does not increase the number of lawn leaving events when animals are not pre-treated with all-trans retinal. Statistics by Wilcoxon rank-sum test. **(H)** Cumulative distribution of time until the first head poke reversal or lawn leaving event while the light is OFF. Statistics by Kolmogorov-Smirnov 2-sample test. **(I-J)** Quantification of roaming and leaving in animals with or without pre-treatment with all-trans retinal without light exposure. (-ATR n = 67, +ATR n = 57). **(I)** Pre-treatment with all-trans retinal does not alter the fraction of time roaming. Statistics by Student’s t-test on logit-transformed data. **(J)** Pre-treatment with all-trans retinal does not alter the number of lawn leaving events per animal. Statistics by Mann-Whitney U test. Statistics: ns not significant (p > 0.05), **** p < 10^-4^ Violin plots show median and interquartile range. In (D,E,G), each dot pair connected by a line represents data from a single animal. In (C), each dot pair connected by a line represents data preceding a single lawn leaving event. See Supplementary Table 3.

**Figure 4 – supplement 1:**
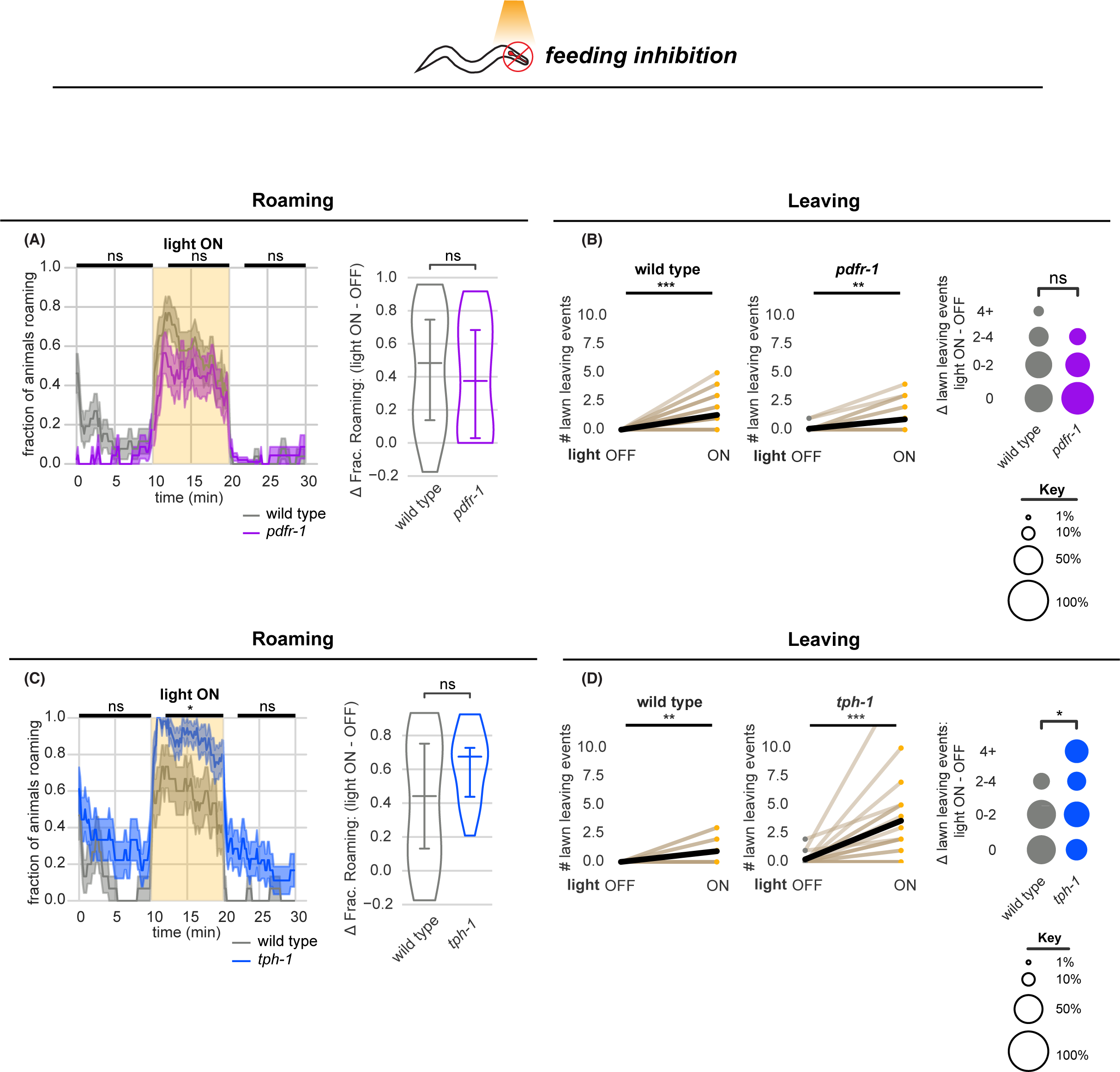
Optogenetic feeding inhibition stimulates roaming and leaving in neuromodulatory mutants. **(A-B)** Optogenetic feeding inhibition in *pdfr-1* mutants. **(A)** Left, Fraction of animals roaming before, during and after optogenetic feeding inhibition in *pdfr-1* mutants and paired wild type controls. Statistics by Student’s t-test comparing *pdfr-1* and matched wild type data averaged and logit-transformed in intervals denoted by black bars above: Data compared at 0-10, 12-20, 22-30 minutes. Right, Difference in fraction of time roaming between light ON and light OFF conditions per animal. Statistics by Mann-Whitney U test. **(B)** Left and Center, *pdfr-1* animals and wild type controls execute more lawn leaving events under feeding inhibition while light is ON. Statistics by Wilcoxon rank-sum test. Right, difference in number of lawn leaving events between light ON and light OFF conditions per animal. Statistics by Mann-Whitney U test. (*pdfr-1* n = 23, wild type controls n = 27) **(C-D)** Optogenetic feeding inhibition in *tph-1* mutants. **(C)** Same as (A) for *tph-1* animals and matched controls. **(D)** Same as (B) for *tph-1* animals and matched controls. (*tph-1* n = 18, wild type controls n = 16) Statistics: ns not significant (p > 0.05), * p < 0.05, ** p < 0.01, *** p < 10^-3^ Violin plots show median and interquartile range. In time-averages, dark line represents the mean and shaded region represents the standard error. In paired plots, each dot pair connected by a line represents data from a single animal. Thick black line indicates the average. See Supplementary Table 4.

**Figure 5 – supplement 1:**
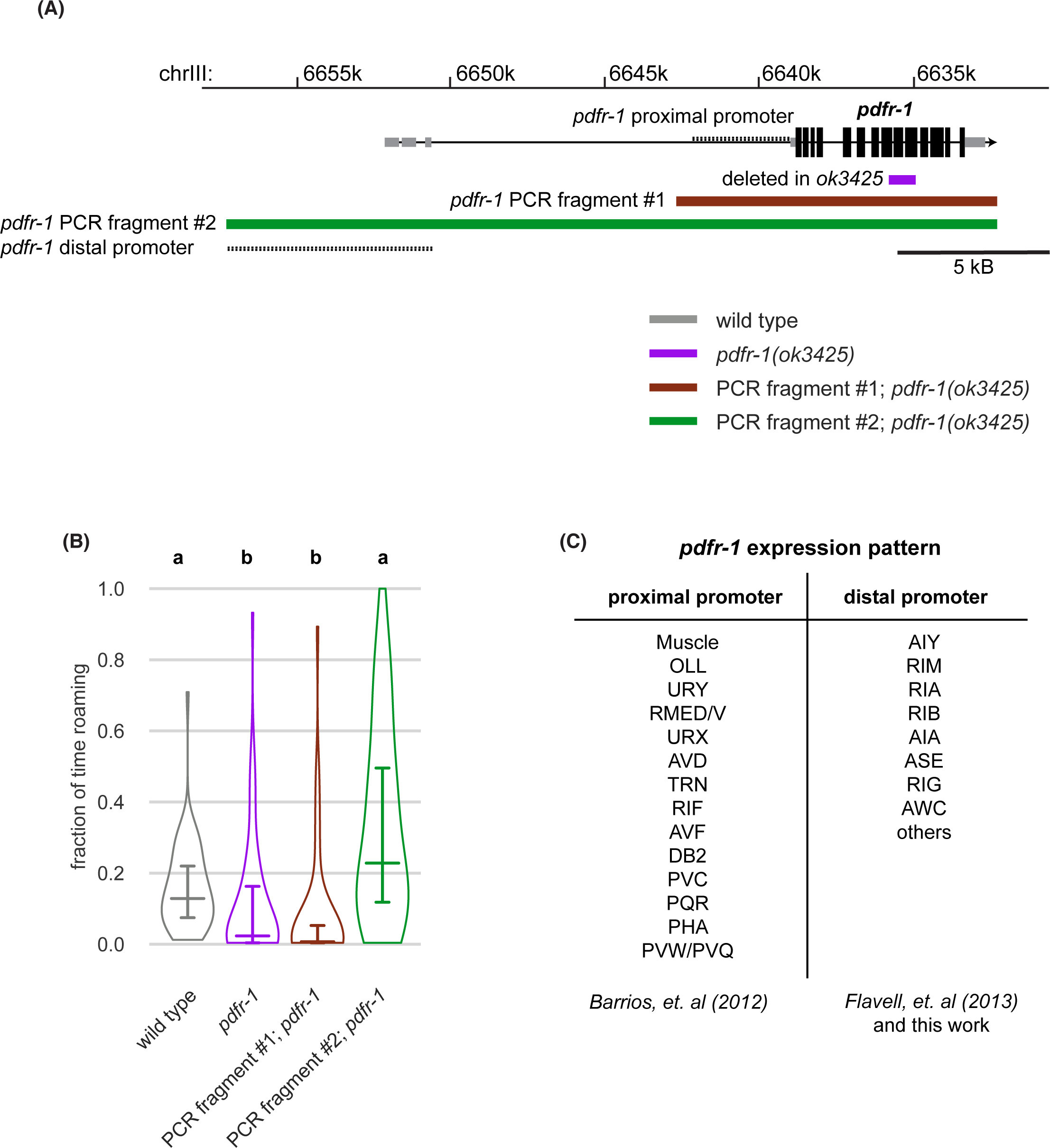
*pdfr-1* genomic characteristics and expression patterns. **(A)** Genomic region surrounding *pdfr-1* (black boxes represent coding exons, gray boxes are non-coding exons), region deleted in *pdfr-1(ok3425)* mutants, proximal and distal promoters, PCR fragments used in transgenic rescue experiments. **(B)** Rescue of *pdfr-1* roaming phenotype by transgenic expression from the PCR fragments illustrated in (A). Different letters mark significant differences in roaming by logit-transformation followed by Tukey’s post hoc test (wild type n = 55, *pdfr-1* n = 59, PCR fragment #1; *pdfr-1* n = 46, PCR fragment #2; *pdfr-1* n = 51). Violin plots show median and interquartile range. **(C)** *pdfr-1* expression patterns from the proximal and distal promoters as reported (Barrios, et al (2012), Flavell, et al (2013)) and from this work. See Supplementary Table 5.

**Figure 5 – supplement 2:**
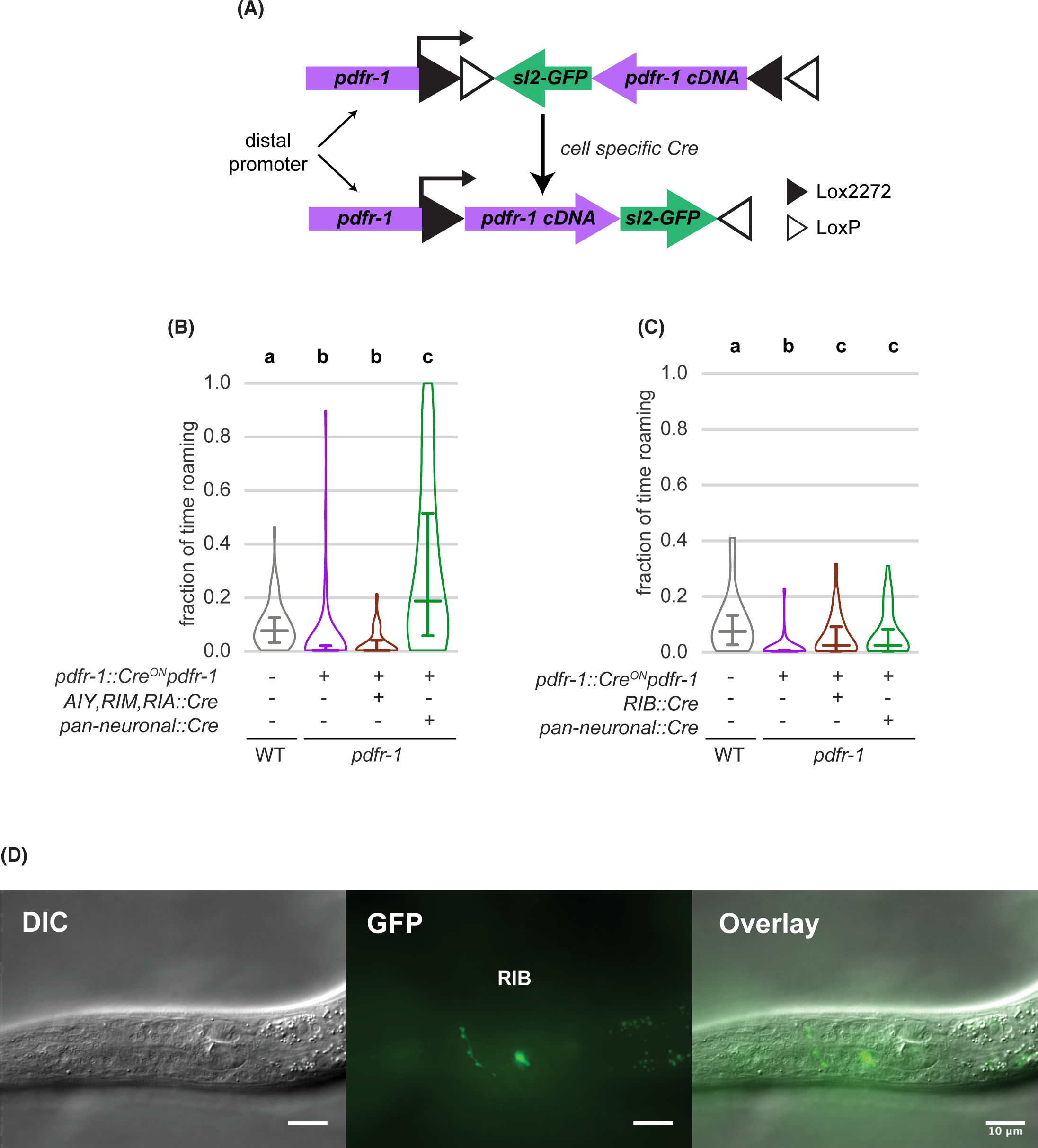
Transgenic rescue of *pdfr-1* in RIB neurons restores roaming. **(A)** Schematic depicting intersectional cell-specific rescue of *pdfr-1* using an inverted Cre-Lox strategy. **(B)** Restoring *pdfr-1* expression in AIY, RIM, and RIA neurons did not rescue roaming on small lawns (wild type n = 64, inverted *pdfr-1* transgene n = 64, AIY, RIM, RIA *pdfr-1* rescue n = 64, pan-neuronal::Cre rescue n = 67). **(C)** Restoring *pdfr-1* expression in RIB neurons rescued roaming comparably to pan-neuronal rescue on small lawns (wild type n = 27, inverted *pdfr-1* transgene n = 32, RIB *pdfr-1* rescue n = 33, pan-neuronal::Cre rescue n = 31). **(D)** Differential interference contrast (DIC), GFP fluorescence and composite images of an animal expressing *pdfr-1* in the RIB neurons using the transgenic strategy shown in (A). Scale bar is 10μm. Statistics: In (B-C), different letters mark significant differences in roaming by logit-transformation followed by Tukey’s post hoc test. Violin plots (B,C) show median and interquartile range. See Supplementary Table 5.

**Figure 5 – supplement 3:**
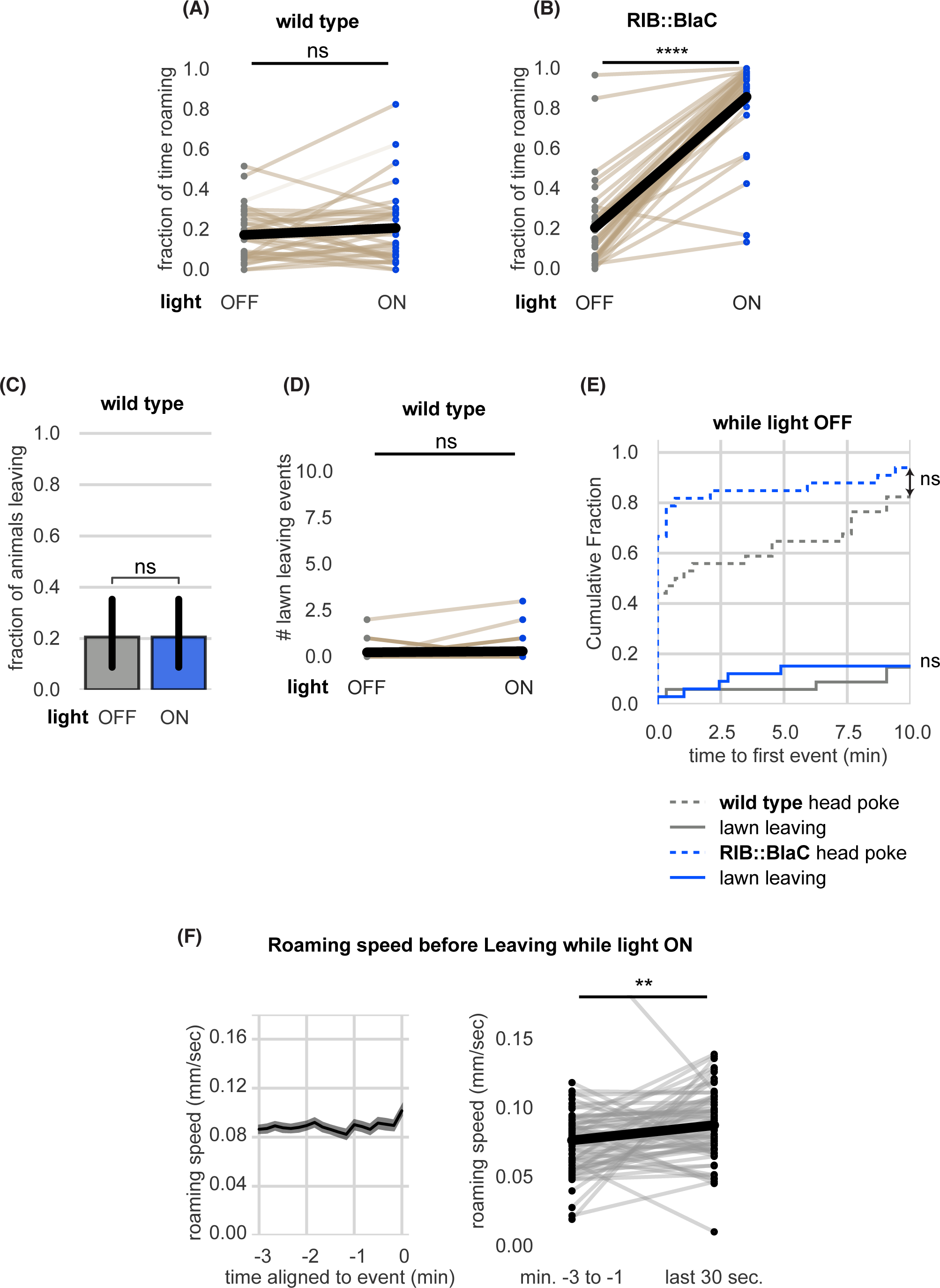
Further quantification and controls of RIB::BlaC experiments. **(A-E)** Quantification of RIB::BlaC controls. **(A)** Blue light stimulation does not induce roaming in wild type animals. Statistics by paired t-test on logit-transformed data. **(B)** Blue light stimulation reliably induces roaming in RIB::BlaC animals. Statistics by paired t-test on logit-transformed data. **(C)** Blue light stimulation does not elevate the fraction of wild type animals that leave lawns. Statistics by Fisher’s exact test. **(D)** Blue light stimulation does not elevate the number of lawn leaving events per wild type animal. Statistics by Wilcoxon rank-sum test. **(E)** Cumulative distribution of time until the first head poke reversal or lawn leaving event while the light is OFF. Statistics by Kolmogorov-Smirnov 2-sample test. **(F)** Roaming animals only accelerate slightly before leaving during RIB BlaC stimulation. Left, mean roaming speed of animals before leaving. Right, quantification of roaming speed changes before leaving. Statistics by Wilcoxon rank-sum test. (RIB BlaC n = 35, wild type n = 34). Statistics: ns not significant (p > 0.05), **** p < 10^-4^ In paired plots (A,B,D), each dot pair connected by a line represents data from a single animal. Thick black line indicates the average. See Supplementary Table 5.

**Figure 6 – supplement 1:**
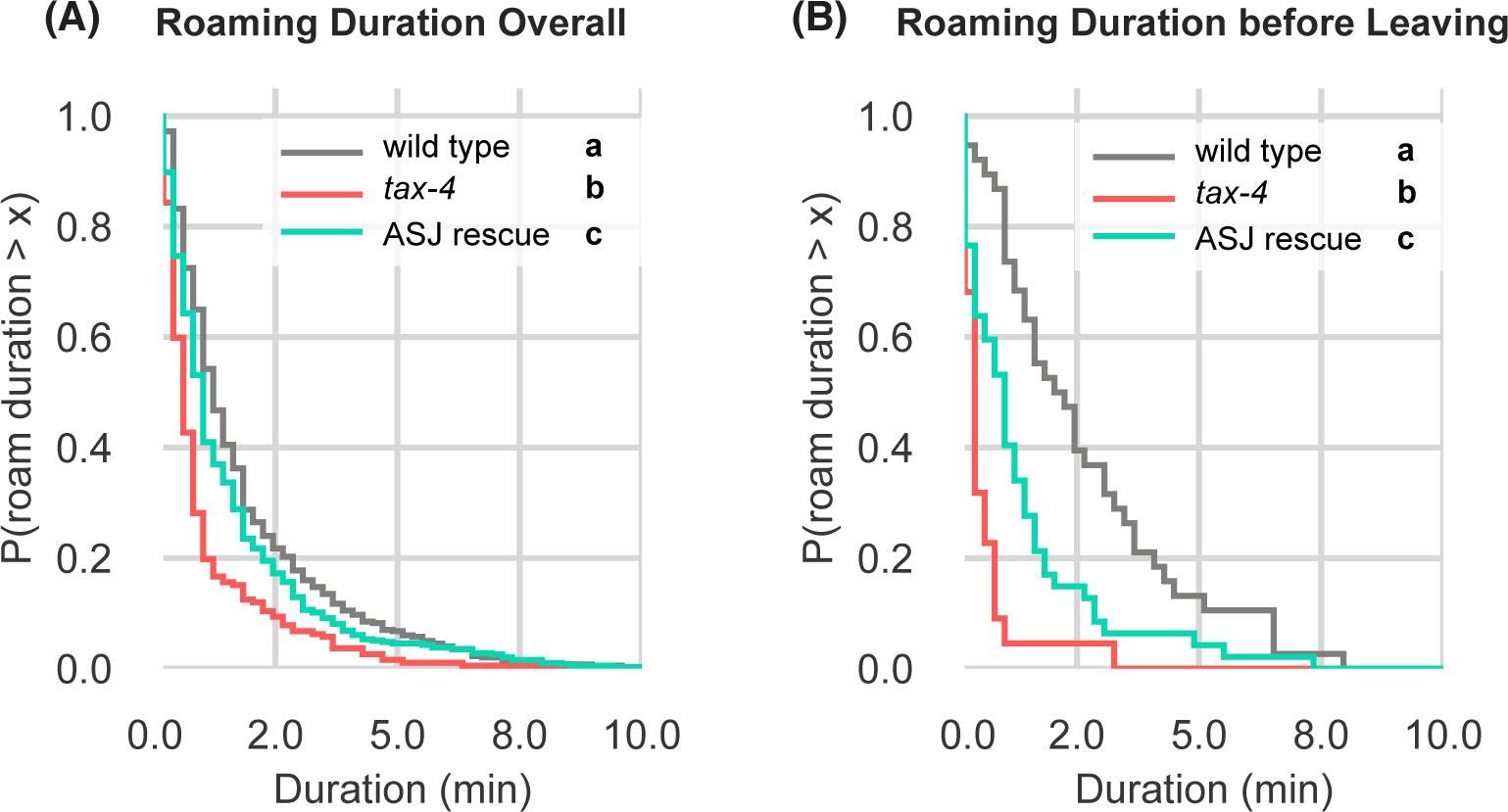
Further quantification of *tax-4* mutants and *tax-4* ASJ rescue on edible food **(A)** Complementary cumulative distribution functions (ccdfs) for overall roam state durations for wild type, *tax-4*, and *tax-4* rescued in ASJ. Different letters mark significant differences by Kolmogorov-Smirnov 2-sample tests with Bonferroni correction. **(B)** Same as (A) for roam state durations before leaving events. Wild type n = 143, *tax-4* n = 156, *tax-4* ASJ rescue n = 148. Statistics: different letters indicate significant differences. See Supplementary Table 6.

**Figure 6 – supplement 2:**
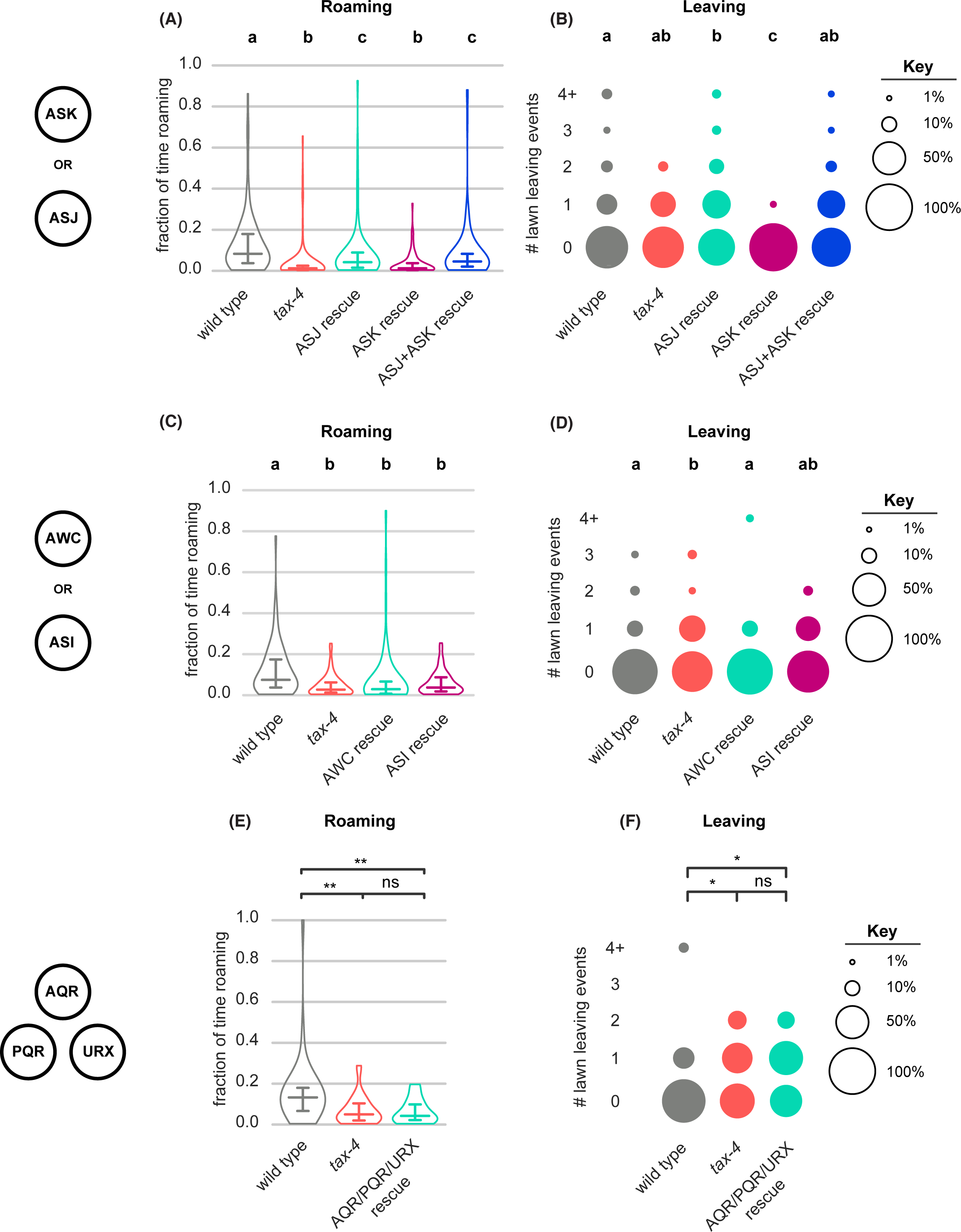
*tax-4* rescue in additional *tax-4-*expressing neurons **(A-B)** Quantification of roaming and leaving in *tax-4* mutants with *tax-4* rescued in ASJ, ASK or ASJ+ASK neurons (wild type n = 95, *tax-4* n = 104, ASJ rescue n = 91, ASK rescue n = 101, ASJ+ASK rescue n = 105). **(A)** ASJ or ASJ+ASK *tax-4* rescue restores roaming to *tax-4* mutants. **(B)** ASK *tax-4* rescue suppresses lawn leaving below *tax-4* levels. **(C-D)** Quantification of roaming and leaving in *tax-4* mutants with *tax-4* rescued in AWC or ASI neurons (wild type n = 79, *tax-4* n = 88, AWC rescue n = 68, ASI rescue n = 77). **(C)** AWC or ASI *tax-4* rescue fails to restore roaming to *tax-4* mutants. **(D)** AWC *tax-4* rescue restores lawn leaving to wild-type levels. **(E-F)** Quantification of roaming and leaving in *tax-4* mutants with *tax-4* rescued in AQR/PQR/URX neurons (wild type n = 35, *tax-4* n = 32, AQR/PQR/URX rescue n = 19). **(E)** AQR/PQR/URX *tax-4* rescue fails to restore roaming to *tax-4* mutants. **(F)** AQR/PQR/URX *tax-4* rescue fails to restore lawn leaving. Violin plots (A,C,E) show median and interquartile range. Statistics: Different letters mark significant differences. ns not significant (p > 0.05), * p < 0.05, ** p < 0.01, *** p < 10^-3^. Statistical tests for differences in fraction of time roaming by one-way ANOVA followed by Tukey’s post hoc test on logit-transformed data. Statistical tests for differences in lawn leaving by Kruskal-Wallis test followed by Dunn’s multiple comparisons test. See Supplementary Table 6.

**Figure 6 – supplement 3:**
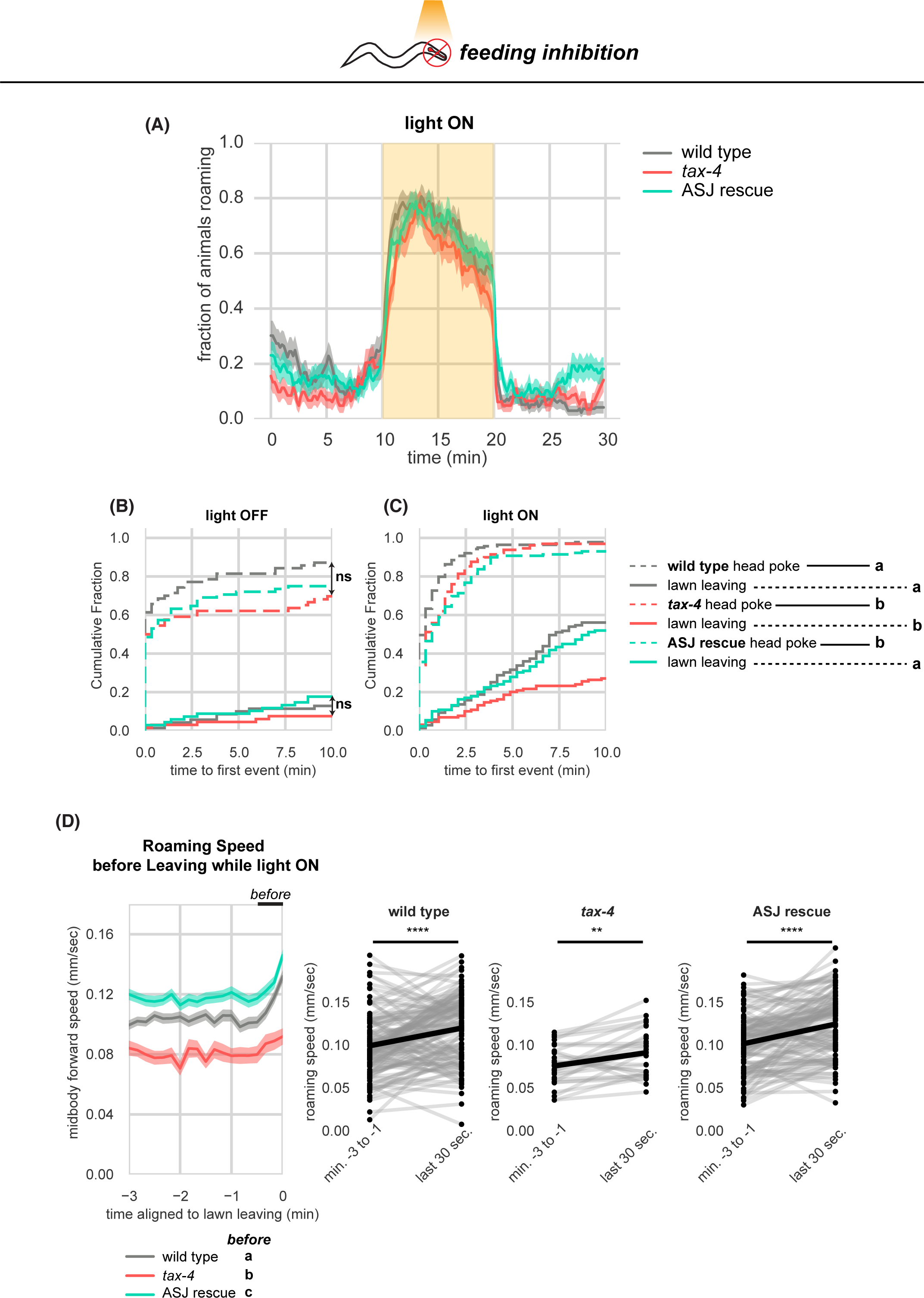
Further quantification of *tax-4* mutants and *tax-4* rescue upon optogenetic feeding inhibition **(A-D)** Effects of *tax-4* on roaming and leaving behavior induced by optogenetic feeding inhibition. **(A)** Fraction of animals roaming before, during, and after optogenetic feeding inhibition. Light ON period denoted by yellow shading. **(B)** Cumulative distribution of time until the first head poke reversal or lawn leaving event while the light is OFF. Statistics by all pairwise Kolmogorov-Smirnov 2-sample tests, Bonferroni corrected. **(C)** Same as (D) for when the light is ON. Pairwise significant differences in time to first lawn leaving are indicated in the figure legend. **(D)** Left, Roaming speed before leaving while the light is ON computed at times when less than 10% of aligned traces had missing data. Roaming speed from the last 30 seconds is averaged and compared by statistical testing: significant differences are reported in the figure legend under “before”. Right, paired plots indicate the average speed from minutes −3 to −1 and in the last 30 seconds before leaving per animal. Each dot pair connected by a line represents data preceding a single lawn leaving event. Thick black line indicates the average. Statistics: ** p < 0.01, *** p < 10^-3^, Different letters mark significant differences. Violin plots (B,C) show median and interquartile range. See Supplementary Table 6.

**Figure 6 – supplement 4:**
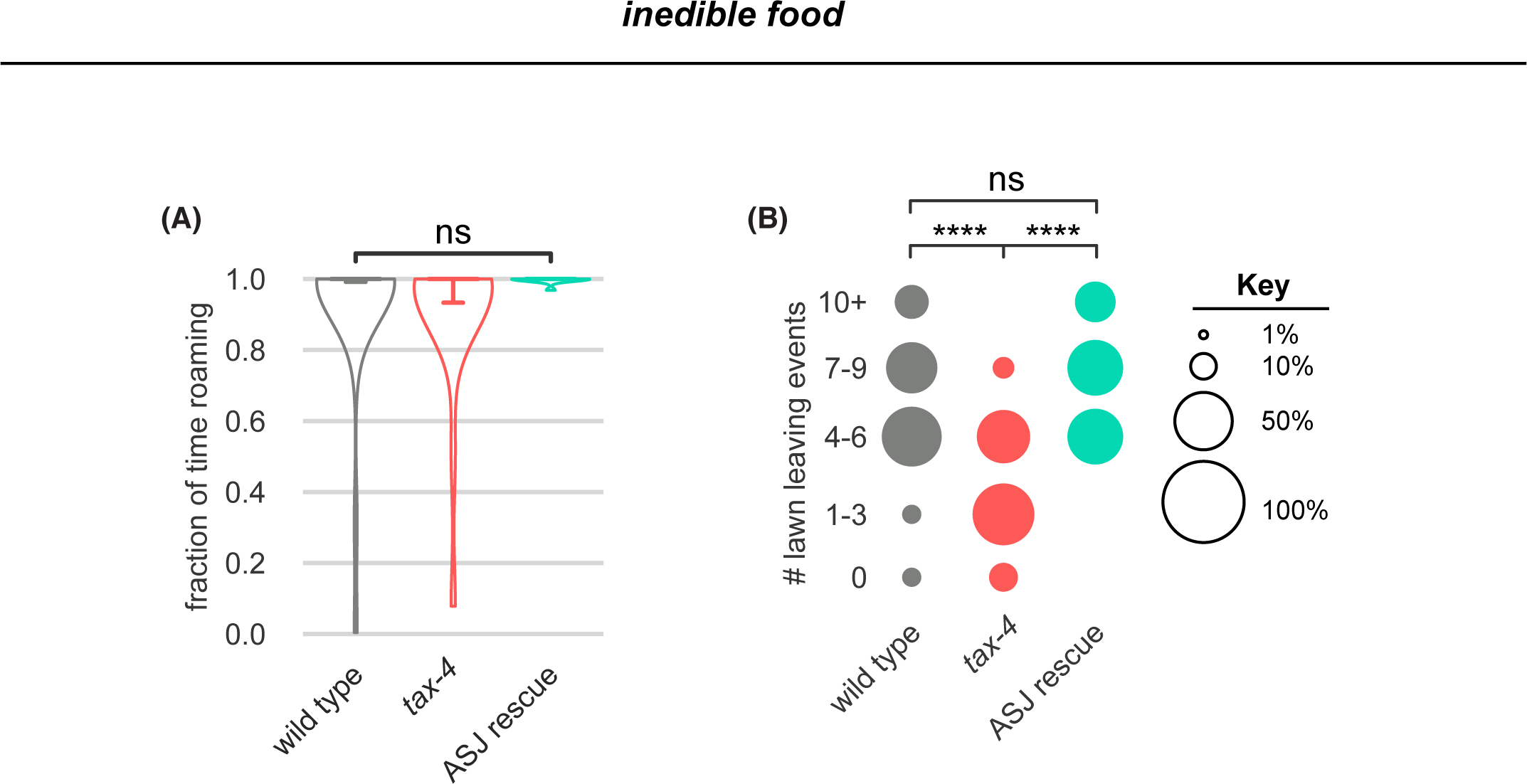
*tax-4* mutants leave less on inedible food and ASJ rescue restores leaving. **(A-B)** In these panels, wild type n = 30, *tax-4* n = 22, *tax-4* ASJ rescue n = 10 on inedible food generated by pre-treatment of bacterial lawns with aztreonam before growth. **(A)** Wild type, *tax-4* and *tax-4* ASJ rescue animals all roam near-constantly on inedible food. Statistical tests of differences in roaming by one-way ANOVA followed by Tukey’s post hoc test on logit-transformed data. **(B)** *tax-4* mutants leave lawns of inedible bacteria less frequently than wild type. *tax-4* ASJ rescue restores lawn leaving. Statistical tests of differences in lawn leaving by Kruskal-Wallis followed by Dunn’s multiple comparisons test. See Supplementary Table 6.

## REFERENCES

Asahina, K., Watanabe, K., Duistermars, B. J., Hoopfer, E., González, C. R., Eyjólfsdóttir, E. A., Perona, P., & Anderson, D. J. (2014). Tachykinin-expressing neurons control male-specific aggressive arousal in *Drosophila*. Cell, 156(1–2), 221–235. 10.1016/j.cell.2013.11.045

Ayala-Orozco, B., Cocho, G., Larralde, H., Ramos-Fernandez, G., Mateos, J. L., & Miramontes, O. (2004). Lévy walk patterns in the foraging movements of spider monkeys (*Ateles geoffroyi*). Behavioral Ecology and Sociobiology, 55(3), 223–230. 10.1007/s00265-003-0700-6

Bargmann, C. Chemosensation in *C. elegans*. (October 22, 2006) WormBook. ed. The *C. elegans* Research Community, WormBook, 10.1895/wormbook.1.123.1, http://www.wormbook.org.

Ben Arous, J., Laffont, S., & Chatenay, D. (2009). Molecular and sensory basis of a food related two-state behavior in *C. elegans*. PLoS ONE, 4(10), e7584. 10.1371/journal.pone.0007584

Bendesky, A., Tsunozaki, M., Rockman, M. V., Kruglyak, L., & Bargmann, C. I. (2011). Catecholamine receptor polymorphisms affect decision-making in *C. elegans*. Nature, 472(7343), 313–318. 10.1038/nature09821

Berman, G. J., Choi, D. M., Bialek, W., & Shaevitz, J. W. (2014). Mapping the stereotyped behaviour of freely moving fruit flies. Journal of the Royal Society Interface, 11(99). 10.1098/rsif.2014.0672

Buchanan, E. K., Linderman, S., & Paninski, L. (2017). Quantifying the behavioral dynamics of *C. elegans* with autoregressive hidden Markov models. NIPS, 1–5. https://openreview.net/pdf?id=o_a_bBItkeO

Cermak, N., Yu, S. K., Clark, R., Huang, Y. C., Baskoylu, S. N., & Flavell, S. W. (2020). Whole-organism behavioral profiling reveals a role for dopamine in state dependent motor program coupling in *C. elegans*. ELife, 9, 1–34. 10.7554/eLife.57093

Charnov, E. L. (1976). Optimal foraging, the marginal value theorem. Theoretical Population Biology, 9(2), 129–136. 10.1016/0040-5809(76)90040-X

Cheung, B. H. H., Cohen, M., Rogers, C., Albayram, O., & De Bono, M. (2005). Experience-dependent modulation of *C. elegans* behavior by ambient oxygen. Current Biology, 15(10), 905–917. 10.1016/j.cub.2005.04.017

Coen, P., Clemens, J., Weinstein, A. J., Pacheco, D. A., Deng, Y., & Murthy, M. (2014). Dynamic sensory cues shape song structure in *Drosophila*. Nature, 507(7491), 233–237. 10.1038/nature13131

Dal Bello, M., Pérez-Escudero, A., Schroeder, F. C., & Gore, J. (2021). Inversion of pheromone preference optimizes foraging in *C. elegans*. ELife, 10, 1–21. 10.7554/eLife.58144

Datta, S. R., Anderson, D. J., Branson, K., Perona, P., & Leifer, A. (2019). Computational neuroethology: A call to action. Neuron, 104(1), 11–24. 10.1016/j.neuron.2019.09.038

Davis, K., Mitchell, C., Weissenfels, O., Bai, J., Raizen, D. M., Ailion, M., & Topalidou, I. (2023). G protein-coupled receptor kinase-2 (GRK-2) controls exploration through neuropeptide signaling in *Caenorhabditis elegans*. PLOS Genetics, 19(1), e1010613. 10.1371/journal.pgen.1010613

Faul, F., Erdfelder, E., Buchner, A., & Lang, A.-G. (2009). Statistical power analyses using G*Power 3.1: Tests for correlation and regression analyses. Behavior Research Methods, 41, 1149–1160.

Flavell, S. W., Gogolla, N., Lovett-Barron, M., & Zelikowsky, M. (2022). The emergence and influence of internal states. Neuron, 110(16), 2545–2570. 10.1016/j.neuron.2022.04.030

Flavell, S. W., Pokala, N., Macosko, E. Z., Albrecht, D. R., Larsch, J., & Bargmann, C. I. (2013). Serotonin and the neuropeptide PDF initiate and extend opposing behavioral states in *C. elegans*. Cell, 154(5), 1023–1035. 10.1016/j.cell.2013.08.001

Fujiwara, M., Sengupta, P., & McIntire, S. L. (2002). Regulation of body size and behavioral state of *C. elegans* by sensory perception and the *egl-4* cGMP-dependent protein kinase. Neuron, 36(6), 1091–1102. 10.1016/S0896-6273(02)01093-0

Gallagher, T., Bjorness, T., Greene, R., You, Y.-J., & Avery, L. (2013). The geometry of locomotive behavioral states in *C. elegans*. PLoS ONE, 8(3), e59865. 10.1371/journal.pone.0059865

Gold, J. I., & Shadlen, M. N. (2007). The neural basis of decision making. Annual Review of Neuroscience, 30(1), 535–574. 10.1146/annurev.neuro.29.051605.113038

Greene, J. S., Dobosiewicz, M., Butcher, R. A., McGrath, P. T., & Bargmann, C. I. (2016). Regulatory changes in two chemoreceptor genes contribute to a *Caenorhabditis elegans* QTL for foraging behavior. ELife, 5, 1–19. 10.7554/eLife.21454

Gruninger, T. R., Gualberto, D. G., & Garcia, L. R. (2008). Sensory perception of food and insulin-like signals influence seizure susceptibility. PLoS Genetics, 4(7). 10.1371/journal.pgen.1000117

Hapiak, V., Summers, P., Ortega, A., Law, W. J., Stein, A., & Komuniecki, R. (2013). Neuropeptides amplify and focus the monoaminergic inhibition of nociception in *Caenorhabditis elegans*. Journal of Neuroscience, 33(35), 14107–14116. 10.1523/JNEUROSCI.1324-13.2013

Hattar, S., Liao, H. W., Takao, M., Berson, D. M., & Yau, K. W. (2002). Melanopsin-containing retinal ganglion cells: Architecture, projections, and intrinsic photosensitivity. Science, 295(5557), 1065–1070. 10.1126/science.1069609

Hill, A. J., Mansfield, R., Lopez, J. M. N. G., Raizen, D. M., & Van Buskirk, C. (2014). Cellular stress induces a protective sleep-like state in *C. elegans*. Current Biology, 24(20), 2399–2405. 10.1016/j.cub.2014.08.040

Hindmarsh Sten, T., Li, R., Otopalik, A., & Ruta, V. (2021). Sexual arousal gates visual processing during *Drosophila* courtship. Nature, 595(7868), 549–553. 10.1038/s41586-021-03714-w

Hong, W., Kim, D. W., & Anderson, D. J. (2014). Antagonistic control of social versus repetitive self-grooming behaviors by separable amygdala neuronal subsets. Cell, 158(6), 1348–1361. 10.1016/j.cell.2014.07.049

Janssen, T., Husson, S. J., Lindemans, M., Mertens, I., Rademakers, S., Donck, K. Ver, Geysen, J., Jansen, G., & Schoofs, L. (2008). Functional characterization of three G protein-coupled receptors for Pigment Dispersing Factors in *Caenorhabditis elegans*. Journal of Biological Chemistry, 283(22), 15241–15249. 10.1074/jbc.M709060200

Jang, H., Levy, S., Flavell, S. W., Mende, F., Latham, R., Zimmer, M., & Bargmann, C. I. (2017). Dissection of neuronal gap junction circuits that regulate social behavior in *Caenorhabditis elegans*. Proceedings of the National Academy of Sciences USA, 114(7), E1263–E1272. 10.1073/pnas.1621274114

Javer, A., Ripoll-Sánchez, L., & Brown, A. E. X. (2018). Powerful and interpretable behavioural features for quantitative phenotyping of *Caenorhabditis elegans*. Philosophical Transactions of the Royal Society B: Biological Sciences. 10.1098/rstb.2017.0375

Ji, N., Madan, G. K., Fabre, G. I., Dayan, A., Baker, C. M., Kramer, T. S., Nwabudike, I., & Flavell, S. W. (2021). A neural circuit for flexible control of persistent behavioral states. ELife, 10. 10.7554/eLife.62889

Kato, S., Kaplan, H. S., Schrödel, T., Skora, S., Lindsay, T. H., Yemini, E., Lockery, S., & Zimmer, M. (2015). Global brain dynamics embed the motor command sequence of *Caenorhabditis elegans*. Cell, 163(3), 656–669. 10.1016/j.cell.2015.09.034

Kennedy, A., Asahina, K., Hoopfer, E., Inagaki, H., Jung, Y., Lee, H., Remedios, R., & Anderson, D. J. (2014). Internal states and behavioral decision-making: Toward an integration of emotion and cognition. Cold Spring Harbor Symposia on Quantitative Biology, 79, 199–210. 10.1101/sqb.2014.79.024984

Lin, J. Y., Knutsen, P. M., Muller, A., Kleinfeld, D., & Tsien, R. Y. (2013). ReaChR: A red-shifted variant of channelrhodopsin enables deep transcranial optogenetic excitation. Nature Neuroscience, 16(10), 1499– 1508. 10.1038/nn.3502

Linderman, S., Antin, B., Zoltowski, D., & Glaser, J. (2020). SSM: Bayesian Learning and Inference for State Space Models (0.0.1). https://github.com/lindermanlab/ssm

Marques, J. C., Li, M., Schaak, D., Robson, D. N., & Li, J. M. (2020). Internal state dynamics shape brainwide activity and foraging behaviour. Nature, 577(7789), 239–243. 10.1038/s41586-019-1858-z

McCloskey, R. J., Fouad, A. D., Churgin, M. A., & Fang-Yen, C. (2017). Food responsiveness regulates episodic behavioral states in *Caenorhabditis elegans*. Journal of Neurophysiology, 117(5), 1911–1934. 10.1152/jn.00555.2016

Meelkop, E., Temmerman, L., Janssen, T., Suetens, N., Beets, I., Van Rompay, L., Shanmugam, N., Husson, S. J., & Schoofs, L. (2012). PDF receptor signaling in *Caenorhabditis elegans* modulates locomotion and egg-laying. Molecular and Cellular Endocrinology, 361(1–2), 232–240. 10.1016/j.mce.2012.05.001

Meisel, J. D., Panda, O., Mahanti, P., Schroeder, F. C., & Kim, D. H. (2014). Chemosensation of bacterial secondary metabolites modulates neuroendocrine signaling and behavior of *C. elegans*. Cell, 159(2), 267–280. 10.1016/j.cell.2014.09.011

Melo, J. A., & Ruvkun, G. (2012). Inactivation of conserved *C. elegans* genes engages pathogen- and xenobiotic-associated defenses. Cell, 149(2), 452–466. 10.1016/j.cell.2012.02.050

Milward, K., Busch, K. E., Murphy, R. J., De Bono, M., & Olofsson, B. (2011). Neuronal and molecular substrates for optimal foraging in *Caenorhabditis elegans*. Proceedings of the National Academy of Sciences USA, 108(51), 20672–20677. 10.1073/pnas.1106134109

Moy, K., Li, W., Tran, H. P., Simonis, V., Story, E., Brandon, C., Furst, J., Raicu, D., & Kim, H. (2015). Computational methods for tracking, quantitative assessment, and visualization of *C. elegans* locomotory behavior. PLoS ONE, 10(12), 1–22. 10.1371/journal.pone.0145870

O’Connell, L. A., Rigney, M. M., Dykstra, D. W., & Hofmann, H. A. (2013). Neuroendocrine mechanisms underlying sensory integration of social signals. Journal of Neuroendocrinology, 25(7), 644–654. 10.1111/jne.12045

O’Donnell, M. P., Fox, B. W., Chao, P. H., Schroeder, F. C., & Sengupta, P. (2020). A neurotransmitter produced by gut bacteria modulates host sensory behaviour. Nature, 583(7816), 415–420. 10.1038/s41586-020-2395-5

Olofsson, B. (2014). The olfactory neuron AWC promotes avoidance of normally palatable food following chronic dietary restriction. Journal of Experimental Biology, 217(10), 1790–1798. 10.1242/jeb.099929

Omura, D. T., Clark, D. A., Samuel, A. D. T., & Horvitz, H. R. (2012). Dopamine signaling is essential for precise rates of locomotion by *C. elegans*. PLoS ONE, 7(6). 10.1371/journal.pone.0038649

Pan, Y., Meissner, G. W., & Baker, B. S. (2012). Joint control of *Drosophila* male courtship behavior by motion cues and activation of male-specific P1 neurons. Proceedings of the National Academy of Sciences USA, 109(25), 10065–10070. 10.1073/pnas.1207107109

Pradel, E., Zhang, Y., Pujol, N., Matsuyama, T., Bargmann, C. I., & Ewbank, J. J. (2007). Detection and avoidance of a natural product from the pathogenic bacterium *Serratia marcescens* by *Caenorhabditis elegans*. Proceedings of the National Academy of Sciences USA, 104(7), 2295–2300. 10.1073/pnas.0610281104

Proekt, A., Banavar, J. R., Maritan, A., & Pfaff, D. W. (2012). Scale invariance in the dynamics of spontaneous behavior. Proceedings of the National Academy of Sciences USA, 109(26), 10564–10569. 10.1073/pnas.1206894109

Pujol, N., Link, E. M., Liu, L. X., Kurz, C. L., Alloing, G., Tan, M. W., Ray, K. P., Solari, R., Johnson, C. D., & Ewbank, J. J. (2001). A reverse genetic analysis of components of the Toll signaling pathway in *Caenorhabditis elegans*. Current Biology, 11(11), 809–821. 10.1016/S0960-9822(01)00241-X

Raizen, D. M., Zimmerman, J. E., Maycock, M. H., Ta, U. D., You, Y. J., Sundaram, M. V., & Pack, A. I. (2008). Lethargus is a *Caenorhabditis elegans* sleep-like state. Nature, 451(7178), 569–572. 10.1038/nature06535

Rhoades, J. L., Nelson, J. C., Nwabudike, I., Yu, S. K., McLachlan, I. G., Madan, G. K., Abebe, E., Powers, J. R., Colón-Ramos, D. A., & Flavell, S. W. (2019). ASICs mediate food responses in an enteric serotonergic neuron that controls foraging behaviors. Cell, 176(1–2), 85–97.e14. 10.1016/j.cell.2018.11.023

Ryu, M. H., Moskvin, O. V., Siltberg-Liberles, J., & Gomelsky, M. (2010). Natural and engineered photoactivated nucleotidyl cyclases for optogenetic applications. Journal of Biological Chemistry, 285(53), 41501–41508. 10.1074/jbc.M110.177600

Sawin, E. R., Ranganathan, R., & Horvitz, H. R. (2000). *C. elegans* locomotory rate is modulated by the environment through a dopaminergic pathway and by experience through a serotonergic pathway. Neuron, 26(3), 619–631. 10.1016/S0896-6273(00)81199-X

Schwarz, R. F., Branicky, R., Grundy, L. J., Schafer, W. R., & Brown, A. E. X. (2015). Changes in postural syntax characterize sensory modulation and natural variation of *C. elegans* Locomotion. PLoS Computational Biology, 11(8), 1–16. 10.1371/journal.pcbi.1004322

Shtonda, B. B., and Avery, L. (2006). Dietary choice behavior in *Caenorhabditis elegans*. Journal of Experimental Biology, 209(1), 89–102. 10.1242/jeb.01955

Stern, S., Kirst, C., & Bargmann, C. I. (2017). Neuromodulatory control of long-term behavioral patterns and individuality across development. Cell, 171(7), 1649–1662.e10. 10.1016/j.cell.2017.10.041

Taghert, P. H., & Nitabach, M. N. (2012). Peptide neuromodulation in invertebrate model systems. Neuron, 76(1), 82–97. 10.1016/j.neuron.2012.08.035

Taylor, S. R., Santpere, G., Weinreb, A., Barrett, A., Reilly, M. B., Xu, C., Varol, E., Oikonomou, P., Glenwinkel, L., McWhirter, R., Poff, A., Basavaraju, M., Rafi, I., Yemini, E., Cook, S. J., Abrams, A., Vidal, B., Cros, C., Tavazoie, S.,… Miller, D. M. (2021). Molecular topography of an entire nervous system. Cell, 184(16), 4329–4347.e23. 10.1016/j.cell.2021.06.023

Tinbergen, N. (1951). The study of instinct. Oxford, Clarendon Press.

Tomar, S. (2006). Converting video formats with FFmpeg. Linux Journal, 2006(146), 10. https://dl.acm.org/doi/fullHtml/10.5555/1134782.1134792

Van Buskirk, C., & Sternberg, P. W. (2007). Epidermal growth factor signaling induces behavioral quiescence in *Caenorhabditis elegans*. Nature Neuroscience, 10(10), 1300–1307. 10.1038/nn1981

Wang, Y., Zhang, X., Xin, Q., Hung, W., Florman, J., Huo, J., Xu, T., Xie, Y., Alkema, M. J., Zhen, M., & Wen, Q. (2020). Flexible motor sequence generation during stereotyped escape responses. ELife. 10.7554/eLife.56942

Weissbourd, B., Ren, J., DeLoach, K. E., Guenthner, C. J., Miyamichi, K., & Luo, L. (2014). Presynaptic partners of dorsal raphe serotonergic and GABAergic neurons. Neuron, 83(3), 645–662. 10.1016/j.neuron.2014.06.024

Wiltschko, A. B., Johnson, M. J., Iurilli, G., Peterson, R. E., Katon, J. M., Pashkovski, S. L., Abraira, V. E., Adams, R. P., & Datta, S. R. (2015). Mapping sub-second structure in mouse behavior. Neuron, 88(6), 1121–1135. 10.1016/j.neuron.2015.11.031

Worthy, S.E., Rojas, G.L., Taylor, C.J., and Glater, E.E. (2018). Identification of odor blend used by *Caenorhabditis elegans* for pathogen recognition. Chem. Senses 43, 169–180. 10.1093/chemse/bjy001

Yoshimura, J., Ichikawa, K., Shoura, M. J., Artiles, K. L., Gabdank, I., Wahba, L., Smith, C. L., Edgley, M. L., Rougvie, A. E., Fire, A. Z., Morishita, S., & Schwarz, E. M. (2019). Recompleting the *Caenorhabditis elegans* genome. Genome Research, 29(6), 1009–1022. 10.1101/gr.244830.118

